# Pyruvate Kinase Activity Regulates Cystine Starvation Induced Ferroptosis through Malic Enzyme 1 in Pancreatic Cancer Cells

**DOI:** 10.1101/2023.09.15.557984

**Authors:** Elliot Ensink, Tessa Jordan, Hyllana C D Medeiros, Galloway Thurston, Anmol Pardal, Lei Yu, Sophia Y. Lunt

## Abstract

Pancreatic ductal adenocarcinoma (PDAC) is an aggressive cancer with high mortality and limited efficacious therapeutic options. PDAC cells undergo metabolic alterations to survive within a nutrient-depleted tumor microenvironment. One critical metabolic shift in PDAC cells occurs through altered isoform expression of the glycolytic enzyme, pyruvate kinase (PK). Pancreatic cancer cells preferentially upregulate pyruvate kinase muscle isoform 2 isoform (PKM2). PKM2 expression reprograms many metabolic pathways, but little is known about its impact on cystine metabolism. Cystine metabolism is critical for supporting survival through its role in defense against ferroptosis, a non-apoptotic iron-dependent form of cell death characterized by unchecked lipid peroxidation. To improve our understanding of the role of PKM2 in cystine metabolism and ferroptosis in PDAC, we generated PKM2 knockout (KO) human PDAC cells. Fascinatingly, PKM2KO cells demonstrate a remarkable resistance to cystine starvation mediated ferroptosis. This resistance to ferroptosis is caused by decreased PK activity, rather than an isoform-specific effect. We further utilized stable isotope tracing to evaluate the impact of glucose and glutamine reprogramming in PKM2KO cells. PKM2KO cells depend on glutamine metabolism to support antioxidant defenses against lipid peroxidation, primarily by increased glutamine flux through the malate aspartate shuttle and utilization of ME1 to produce NADPH. Ferroptosis can be synergistically induced by the combination of PKM2 activation and inhibition of the cystine/glutamate antiporter *in vitro*. Proof-of-concept *in vivo* experiments demonstrate the efficacy of this mechanism as a novel treatment strategy for PDAC.

**Graphical Abstract:** 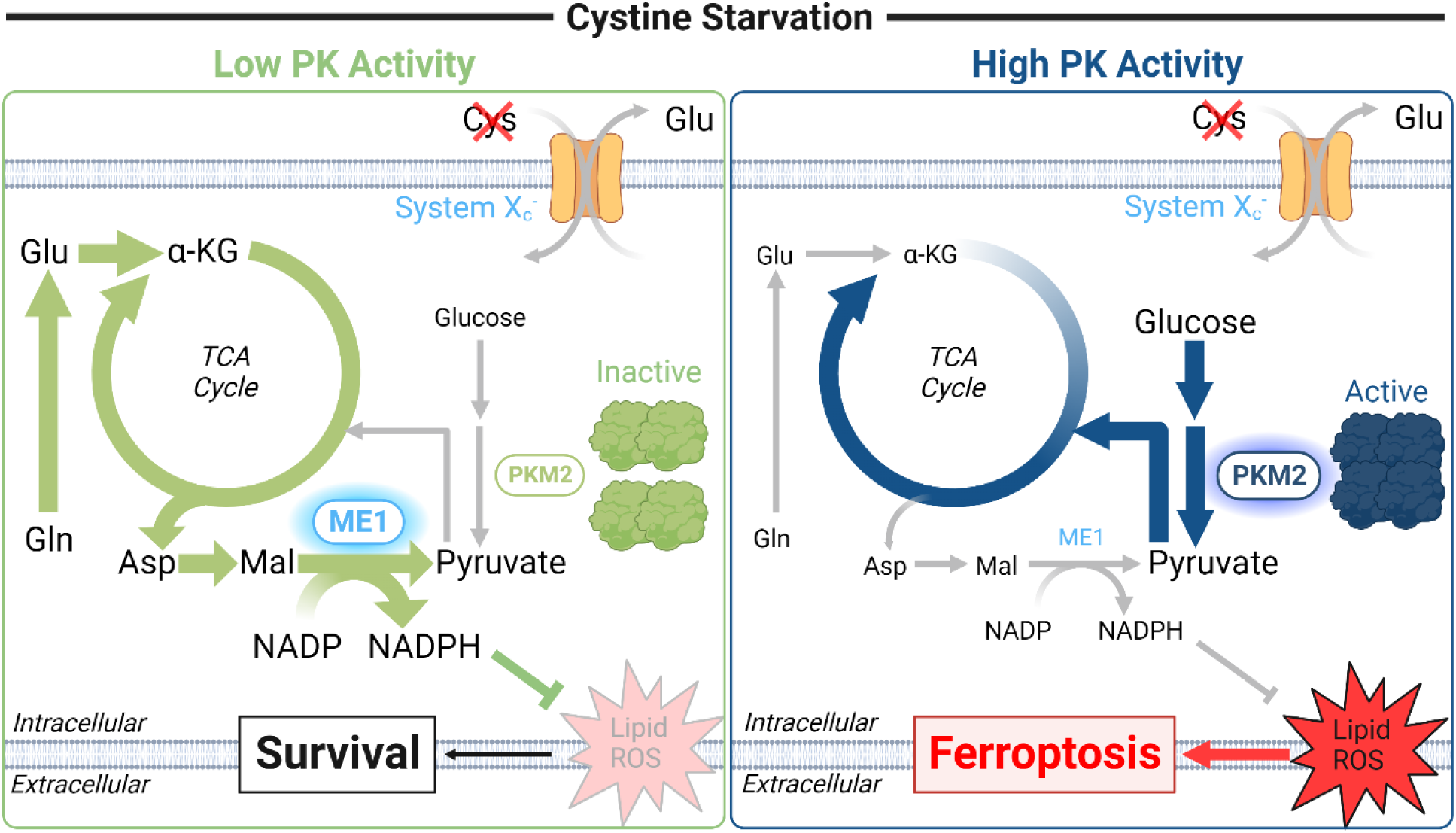

**Highlights:**

- PKM2KO in pancreatic ductal adenocarcinoma (PDAC) cells produces enhanced defense against cystine starvation induced ferroptosis.
- Pharmacologic activation of pyruvate kinase (PK) activity promotes ferroptosis under cystine starvation, while inhibition promotes ferroptosis survival in PDAC cells.
- Decrease in PK activity reprograms glutamine metabolism to increase use of malic enzyme 1 and promote survival under cystine starvation in PDAC cells.
- Cystine starvation and activation of pyruvate kinase synergistically decreases progression of pancreatic cancer *in vivo*.

## Introduction

Pancreatic ductal adenocarcinoma (PDAC) is a devastating disease with a poor prognosis due to limited diagnostic and therapeutic options. PDAC cells are capable of silently progressing and producing metastatic cells before clinical symptoms present or predictive biomarkers can be detected.^1,2^ Pancreatic cancer is currently the fourth leading cause of cancer deaths in the United States and is projected to become the second leading cause of cancer-related deaths by 2030.^3,4^ Disappointingly, recent advances in combinatorial chemotherapy and targeted immunotherapy have failed to significantly improve pancreatic cancer patient outcomes.^2,5^

Disrupting reprogrammed PDAC metabolism is a promising new strategy to improve treatment and survival of pancreatic cancer patients.^6,7^ In the nutrient starved and hypoxic tumor microenvironment, tumor cells undergo many genetic and cellular alterations to reprogram their metabolism.^8^ One consistent adaptation is differential isoform expression of the glycolytic enzyme pyruvate kinase (PK), which converts phosphoenolpyruvate to pyruvate and produces ATP. PDAC cells preferentially switch from the constitutively active PK muscle 1 isoform (PKM1) to the allosterically regulated PK muscle 2 isoform (PKM2).^9–11^ The shift to PKM2 contributes to increased glucose consumption and lactate secretion even in the presence of oxygen, a phenomenon known as the Warburg effect.^12–15^ Overexpression of PKM2 in pancreatic cancer enables rapid proliferation and survival in low glucose conditions by diverting glucose carbon into biosynthetic precursors and altering glutamine metabolism.^16–20^ Glutamine supplies cancer cells with a critical fuel supply for mitochondrial tricarboxylic acid (TCA) cycle and non-essential amino acid production in a process known as anaplerosis when glucose is limited or utilized for other purposes.^18,21^ The shift in PK isoform expression produces a profound reprogramming of complex networks of metabolic pathways, many of which are not well understood, including cysteine metabolism.

Cysteine is critical for cell survival given its dual role in protein synthesis and defense against reactive oxygen species (ROS) as a precursor for tripeptide glutathione and coenzyme A. The antioxidant role of cysteine is especially consequential in pancreatic cancers, which exhibit increased ROS production.^22,23^ Healthy cells utilize circulating cysteine or cysteine biosynthesis to fulfill these roles, but cancer cells instead depend on exogenous cystine, the oxidized dimer of cysteine.^24^ Cystine is acquired predominantly through the cystine/glutamate antiporter (system X_C_^-^) which is a heterodimer of SLC7A11 (also known as xCT) and SLC3A2, and is overexpressed in many cancer cells.^25^ Depletion of extracellular cystine leads to a loss of intracellular glutathione supply and the accumulation of oxidative damage to membrane lipids.^22–24,26^ Uncontrolled lipid peroxidation propagates by reacting with ferrous iron and producing hydroxyl radicals leading to a form of cell death known as ferroptosis.^26,27^ Ferroptosis is fundamentally a product of aberrant metabolic behavior including changes in central carbon metabolism such as increased mitochondrial glutaminolysis and increased dependence on glucose flux through the pentose phosphate pathway to generate NADPH.^28–30^ The impact of PKM2, which also plays a critical role in metabolic reprograming and the management of oxidative stress, on ferroptosis in pancreatic cancer remains poorly characterized. Previous work in our lab demonstrated that PDAC cells use cysteine catabolism to support pyruvate production during PKM2 knockdown.^31^ Additionally, current evidence suggests PKM2 in its inactive dimeric form reprograms central carbon metabolism to increase supply of precursors for biosynthesis of nucleotides and antioxidant defense.^32–35^ To further elucidate reprogrammed PDAC metabolism, we investigated the mechanisms by which PK activity impact cysteine, glucose, and glutamine metabolism in pancreatic cancer cells under nutrient restricted conditions.

Herein, we demonstrate that PKM2 knockout (PKM2KO) in PDAC cells is associated with increased defense against cystine starvation induced ferroptosis. This effect is mediated by decreased PK activity, reprogramming of glutamine metabolism, and utilization of malic enzyme 1 (ME1) to produce NADPH, a critical reducing agent for protection against ROS. Further, we identify that decreased PK activity promotes survival under cystine starvation while activation of PK leads to increased ferroptosis. Lastly, we demonstrate proof-of-concept that the combination of cystine starvation and PKM2 activation is a novel and efficacious strategy for treating PDAC.

## Results

### PKM2KO enhances PDAC survival during cystine starvation

To evaluate the impact of PKM2 on cysteine metabolism, we generated human PDAC cell lines that lack PKM2 expression. PKM isoform selection occurs through mutually exclusive alternative splicing of the PKM gene. Inclusion of exon 9 leads to PKM1 expression and inclusion of exon 10 leads to PKM2 expression.^9,10^ Human PDAC typically expresses PKM2.^32,36^ We selected 2 human PDAC cell lines, AsPC1 and Panc1, due to their sensitivity to cystine starvation.^22^ Using lentiviral clustered regularly interspaced short palindromic repeats (CRISPR)/caspase 9 (cas9) gene editing technology to target exon 10 we selectively knocked out (KO) PKM2 (Fig. 1A). We characterized the expression of PKM isoforms within targeted cells and obtained successful PKM2KO clones for AsPC1 and Panc1. Deletion of exon 10 led to complete loss of PKM2 expression and low level re-expression of PKM1 in all generated clones (Figs. 1B and S1H-I).

**Figure 1.**
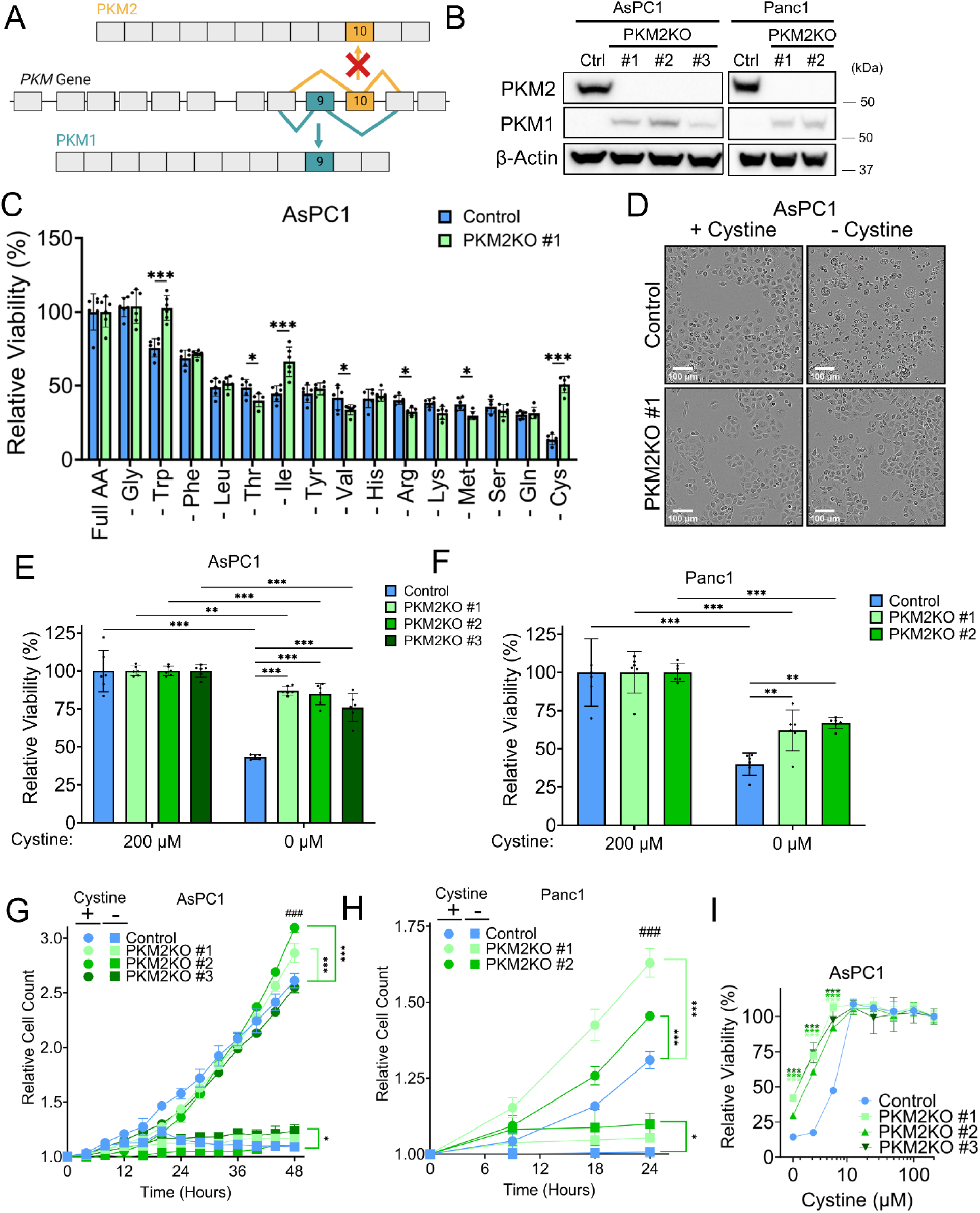
PKM2KO enhances PDAC survival during cystine starvation. **A.** Schematic view of mutually exclusive alternative splicing of *PKM* to produce PKM1 or PKM2. Targeting of exon 10 by CRISPR deletes PKM2 expression. **B.** Western blot of PKM1 and PKM2 in AsPC1 and Panc1 control cells and PKM2KO clones. **C.** Relative viabilities of AsPC1 control and PKM2KO clone #1 in DMEM without each individual amino acid as shown. Significance was assessed by two-way ANOVA and Tukey test. \**p*<0.05, \*\*\**p*<0.001. **D.** Brightfield microscopy images of AsPC1 control and PKM2KO cells under either 200 μM (+) or 0 μM (-) cystine conditions. Scale bar = 100 μm. **E, F.** Relative viabilities of AsPC1 (**E**) and Panc1 (**F**) control and PKM2KO clones under 200 μM or 0 μM cystine. Significance was assessed by two-way ANOVA and Tukey test. \*\**p*<0.01, \*\*\**p*<0.001. **G, H.** Proliferation analysis using Incucyte cell counts of both AsPC1 (**G**) and Panc1 (**H**) control and their respective PKM2KO clones under 200 μM (+) or 0 μM (-) cystine. Significance was assessed by two-way ANOVA and Tukey test. Comparison between Control and PKM2KO cells at endpoint: \**p*<0.05, \*\*\**p*<0.001. Comparison between 200 and 0 μM cystine conditions for each cell line at endpoint: ###*p*<0.001. **I.** Relative viabilities of AsPC1 control and PKM2KO clones under a range of cystine concentrations from 200 μM to 0 μM. Significance was assessed by two-way ANOVA and Dunnet test. \**p*<0.05.

We first evaluated the tolerance of PKM2KO cells to low nutrient stress by assessing the viability of PKM2KO cells in response to depleting each of the individual amino acids typically included in Dulbecco’s Modified Eagle Medium (DMEM). This produced significant differences in survival between controls and PKM2KO clones under starvation of several amino acids (Figs. 1C & S1A-B). The most dramatic and consistent difference was the increased viability of all PKM2KO clones under cystine starvation, the most restrictive condition for the respective control cell lines. Morphologic examination of these cells revealed distinct cell volume shrinkage and membrane blistering in the control cells, characteristic of ferroptosis morphology^37^, and retention of normal morphology in the PKM2KO cells (Figs. 1D and S1E). We then repeated this experiment using DMEM containing physiologic levels of glucose (5 mM) and glutamine (1 mM) and either 200 μM cystine (supraphysiologic) or 0 μM cystine (starvation).^38,39^ Under cystine starvation, the PKM2KO cells had significantly higher viability compared to the PKM2 expressing controls when evaluated using the AlamarBlue viability assay (Fig. 1E-F). Given that the AlamarBlue viability assay relies on metabolic reducing power to determine viability, we also measured viability using trypan blue exclusion, which also showed significantly increased viability of PKM2KO cells under cystine starvation, confirming the effect is not due only to changes in reducing power production (Fig. S1D). To further ensure that the enhanced viability of PKM2KO cells was not simply an artifact of the vector introduced in the control cells, we evaluated the parental wild-type (WT) AsPC1 and Panc1 cells and observed nearly identical sensitivity to cystine starvation as the control cells and significantly lower viability compared to the PKM2KO cells (Fig. S1F-G). We next evaluated the proliferative capacity of the control and PKM2KO cells under cystine starvation. Cystine starvation resulted in dramatic inhibition of cell proliferation in the AsPC1 and Panc1 control and PKM2KO cell lines tested (Fig. 1G-H), indicating that despite the enhanced viability of PKM2KO cells, they are not actively dividing in this environment. We then exposed the AsPC1 PKM2KO cells to a range of cystine concentrations and found that the significant increase in viability in PKM2KO expressing cells occurs as cystine concentrations reach 10 μM or lower (Figs. 1I and S1C). Together, these results suggest that cells that are not expressing PKM2 may represent a sub-population of cells able to persist under stressful low nutrient conditions and allow for tumor survival.^11^

### PKM2KO improves defense against cystine depletion induced ferroptosis in PDAC

Next, we examined the mechanism of cell death occurring in the control cells under cystine starvation. Cystine starvation is known to induce ferroptosis, a non-apoptotic and oxidative damage-related form of cell death driven by iron accumulation and lipid peroxidation.^27^ Under cystine starvation, co-treatment of the AsPC1 and Panc1 cells with the ferroptosis inhibitors ferrostatin-1 (a lipid peroxide inhibitor, FER), trolox (a vitamin E derivative and lipophilic antioxidant; TRO), and deferoxamine (an iron chelator; DFO) significantly restored viability in control cells (Fig. 2A-B) with little effect on the PKM2KO clones. These results indicate that the PKM2 expressing control cells are undergoing ferroptosis while the PKM2KO clones resist this process. Importantly, co-treatment with Z-VAD-FMK (an apoptosis inhibitor; ZVAD) and Necrostatin-1S (a necroptosis inhibitor; NEC) had no significant effect on viability indicating that cell death is not occurring through either apoptosis or necroptosis (Fig. 2A, D). Given that ferroptosis is characterized by an accumulation of unchecked lipid peroxidation, we next measured the ratio of oxidized to reduced lipids using the C11-BODIPY probe, which detects lipid peroxides by shifting from red (∼590 nm) to green (∼510 nm) emission when oxidized. Consistently, PKM2KO cells have nearly no increase in lipid peroxidation compared to the significantly elevated levels in the respective control cells (Figs. 2C-E and S2A). The difference in lipid peroxidation can be eliminated by ferrostatin-1 co-treatment (Fig. 2C-D). AsPC1 and Panc1 PKM2KO cells also have significantly lower general ROS accumulation as measured by the 2’,7’-dichlorodihydrofluorescein diacetate (H2DCFDA) probe, a general detector of ROS (Fig. S2B-C). These observations support the conclusion that PKM2KO in PDAC cells enhances their resistance to cystine starvation induced ferroptosis.

**Figure 2.**
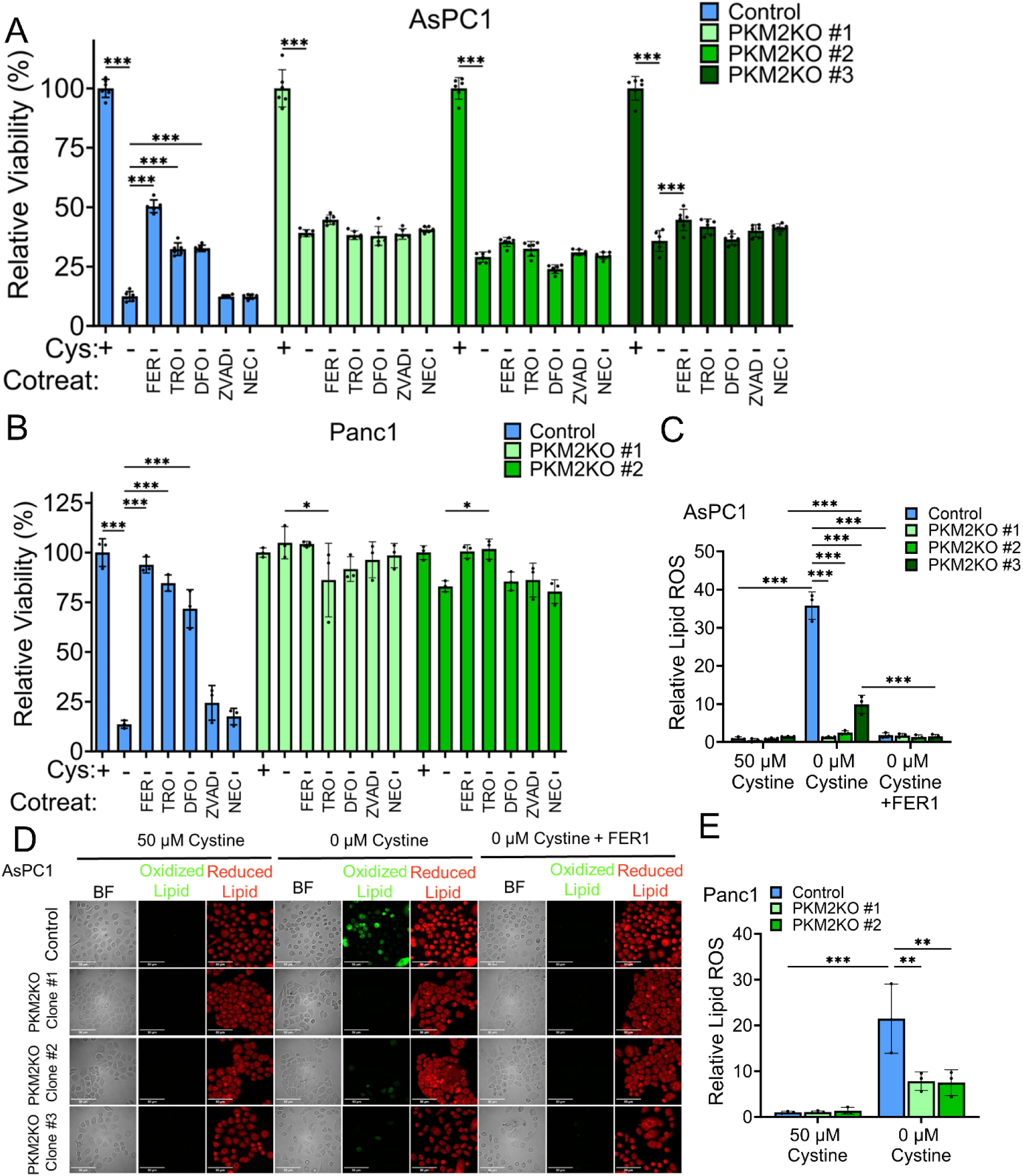
PKM2KO improves defense against cystine depletion induced ferroptosis in PDAC. **A, B.** Relative viabilities of AsPC1 (**A**) and Panc1 (**B**) control and PKM2KO cells under 50 μM cystine (+) and 0 μM cystine (-) co-treated with 5 μM ferrostatin-1 (FER), 100 μM trolox (TRO), 100 μM deferoxamine (DFO), 50 μM Z-VAD-FMK (ZVAD), or 10 μM necrostatin-1S (NEC). Significance by two-way ANOVA. \**p*<0.05, \*\*\**p*<0.001. Multiple hypothesis correction by Tukey test. **C.** Relative lipid peroxidation of AsPC1 control and PKM2KO cells under 50 μM cystine, 0 μM cystine, and 0 μM cystine with 5 μM FER1, visualized by C11-BODIPY. **D.** Representative brightfield and fluorescent images of AsPC1 cell lipid peroxidation quantified in panel C. Scale bar = 50 μm. **E.** Relative lipid peroxidation of Panc1 control and PKM2KO cells under 50 μM cystine and 0 μM cystine, visualized by C11-BODIPY. For **C** and **E,** significance was assessed by two-way ANOVA. \**p*<0.05, \*\*\**p*<0.001. Multiple hypothesis correction by Tukey test.

### Pyruvate kinase activity dictates response to cystine starvation induced ferroptosis

We next sought to identify the specificity of the PK isoform to explain how PKM2KO cells have enhanced survival under cystine starvation. Given that the PKM2KO cells re-express PKM1, it could be either the loss of PKM2 or the gain of PKM1 that is driving the difference in viability. To address these possibilities, we used lentiviral vectors to overexpress PKM1 in the control cells and re-introduced PKM2 expression in the PKM2KO cells for both AsPC1 and Panc1 (Fig. S3A). Evaluation of the response to cystine starvation revealed that neither re-expression of PKM1 nor PKM2 altered the viability response (Fig. S3B-E). However, we observed that the control cells in which PKM1 was overexpressed continued to express PKM2 and similarly the PKM2KO cells with PKM2 re-expressed continue to also express PKM1 at a low level indicating that the re-expression of either PK isoform does not restore cells to their original state (Fig. S3A-B). Thus, the total dosage of PK and activity of PK is likely different with respect to their parental cell lines. Overexpression of PKM1 in the control cells does significantly increase PK activity but did not further inhibit viability likely because they are already maximally inhibited by cystine starvation (Figs. S3F and S3H). PKM2 re-expression significantly increased PK activity under cystine replete conditions, but not under cystine starvation (Figs. S3G and S3I). We then hypothesized that the difference in cystine starvation survival is due to differences in total PK activity, not isoform specificity. We tested our hypothesis by evaluating the activity of PK and found that the AsPC1 PKM2KO cells, despite expressing PKM1 at low levels, had consistently lower overall PK activity regardless of cystine availability, whereas the control cells decreased their PK activity in response to 0 μM cystine (Figs. 3A and S3J), consistent with PKM2’s response to oxidative stress.^40^ We then tested whether modulation of PK activity would influence viability under 0 μM cystine. By treating cells with compound 3k, a potent and specific PKM2 antagonist (Fig. 3B), we were able to dramatically restore viability in four different WT PDAC cell lines (AsPC1, Panc1, MiaPaCa2, and BxPC3) under 0 μM cystine stress (Fig. 3C-F). Additionally, co-treatment of imidazole ketone erastin (IKE)^29^, a potent inhibitor of xCT, with compound 3k restored viability in the majority of WT PDAC cell lines tested (Fig. S3L-O), demonstrating that decreased PK activity can improve resistance to cystine starvation induced ferroptosis. To further confirm our hypothesis, we combined IKE treatment with TEPP-46, a potent and selective PKM2 allosteric agonist (Fig. 3B). The combination treatment synergistically and significantly decreased viability in a concentration dependent manner in each cell line tested except BxPC3 (Fig. 3G-J, K-N). Interestingly, BxPC3 showed strong response to IKE but limited synergistic response when co-treated with TEPP-46 (Fig. 3J, N). Of the 4 WT PDAC cell lines tested, BxPC3 retains residual PKM1 expression suggesting that this cell line likely has reprogrammed glycolytic activity relative to the other WT cells (Fig. S3K). We further evaluated the mechanism of cell death caused by the co-treatment of IKE and TEPP-46 and found that ferrostatin-1 but not Z-VAD-FMK nor necrostatin-1S was able to restore viability, indicating that TEPP-46 is working with IKE to induce ferroptosis and not apoptosis or necroptosis (Fig. S3P-S). Together, these observations present compelling evidence that PK activity dictates the response to cystine starvation induced ferroptosis.

**Figure 3.**
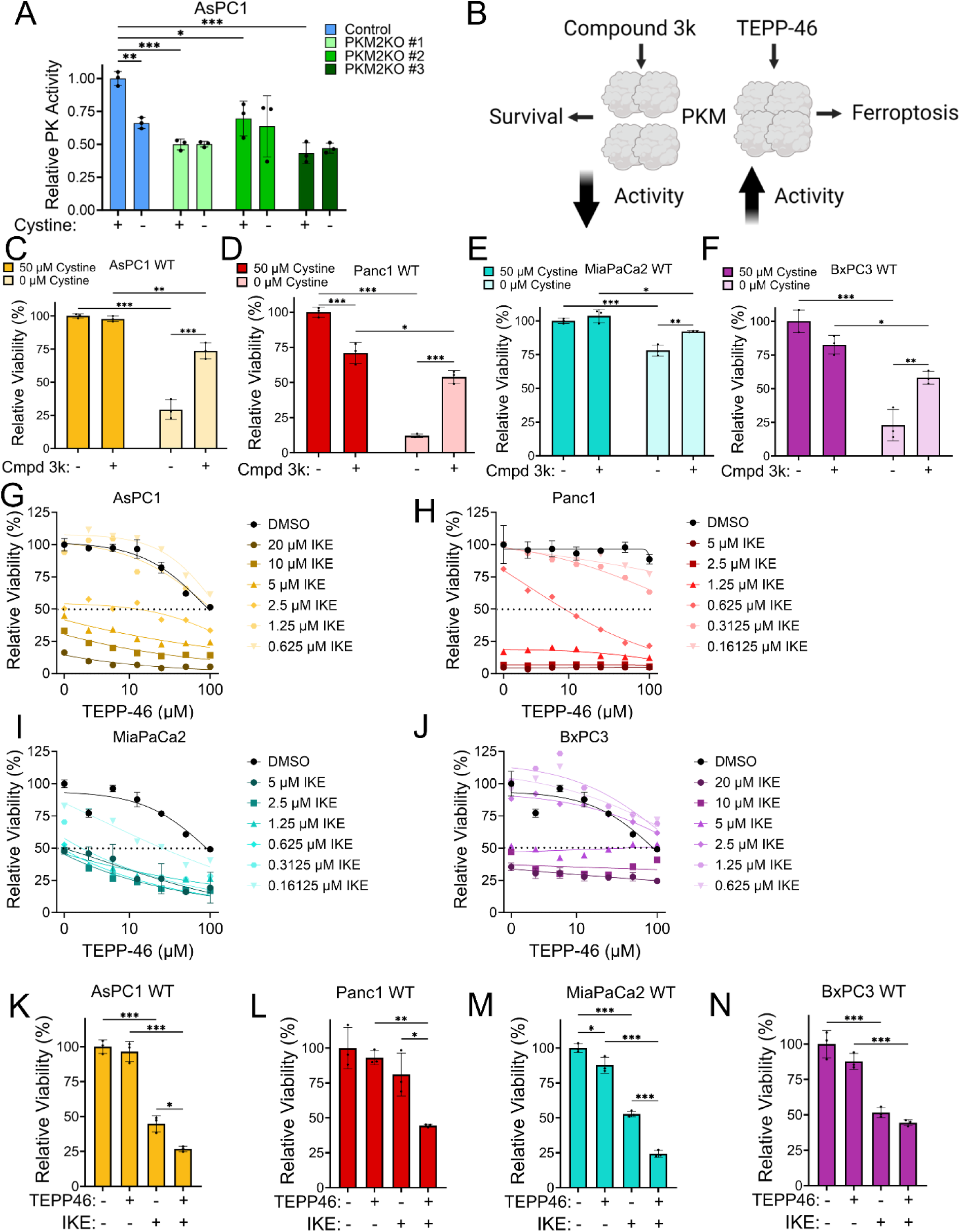
Pyruvate kinase activity dictates response to cystine starvation induced ferroptosis. **A.** Relative PK activity in AsPC1 control and PKM2KO cells under 50 μM (+) and 0 (-) μM cystine. Significance was assessed by two-way ANOVA. \**p*<0.05, \*\*\**p*<0.001. Multiple hypothesis correction by Tukey test. **B.** Schematic showing TEPP-46 promoting the formation of the active tetrameric form of PKM2 and compound 3k inhibiting tetramer formation, producing the less active dimeric form of PKM2. **C-F.** Relative viabilities at 50 μM and 0 μM cystine with (+) or without (-) treatment of 10 μM compound 3k in WT cells: AsPC1 (**C**), Panc1 (**D**), MiaPaCa2 (**E**), and BxPC3 (**F**). Significance was assessed by two-way ANOVA. \**p*<0.05, \*\**p*<0.01, \*\*\**p*<0.001. Multiple hypothesis correction by Tukey test. **G-J.** The effect of IKE and TEPP-46 combination treatment in the range of the indicated concentrations in AsPC1 WT cells (**G**), Panc1 WT cells (**H**), MiaPaCa2 WT cells (**I**), and BxPC3 WT cells (**J**). **K.** Relative viability of AsPC1 WT cells with (+) or without (-) treatment with 5 μM IKE and 12.5 μM TEPP-46. **L.** Relative viability of Panc1 WT cells with (+) or without (-) treatment with 0.625 μM IKE and 12.5 μM TEPP-46. **M.** Relative viability of MiaPaCa2 WT cells with (+) or without (-) treatment with 0.625 μM IKE and 12.5 μM TEPP-46. **N.** Relative viability of BxPC3 WT cells with (+) or without (-) treatment with 5 μM IKE and 12.5 μM TEPP-46. For **L**-**N**, significance was assessed by two-way ANOVA. \**p*<0.05, \*\**p*<0.01, \*\*\**p*<0.001. Multiple hypothesis correction by Tukey test.

### PKM2KO enhances defense against ferroptosis induction specific to cystine starvation

Two key defensive proteins against ferroptosis are xCT and glutathione peroxidase 4 (GPX4). xCT is a component of the cystine/glutamate antiporter and is the transporter by which cancer cells acquire nearly all of their cystine.^41^ GPX4 is primarily responsible for quenching lipid peroxides.^41^ Two of the drugs known to potently induce ferroptosis, IKE and ras selective lethal 3 (RSL3), target xCT and GPX4 respectively (Fig. 4A).^41^ We evaluated the expression of these proteins in our cell lines and found that under cystine starvation, AsPC1 control cells increase xCT to a greater extent than PKM2KO cells, while Panc1 control and PKM2KO cells have similar expression (Figs. 4B and S4F). Both AsPC1 and Panc1 control cells have decreased GPX4 expression under cystine starvation, but PKM2KO cells retain equivocal expression of GPX4 (Fig. 4B and S4F). This suggests that the presence of PKM2 may directly or indirectly modulate expression of these genes through a yet to be determined mechanism.

**Figure 4.**
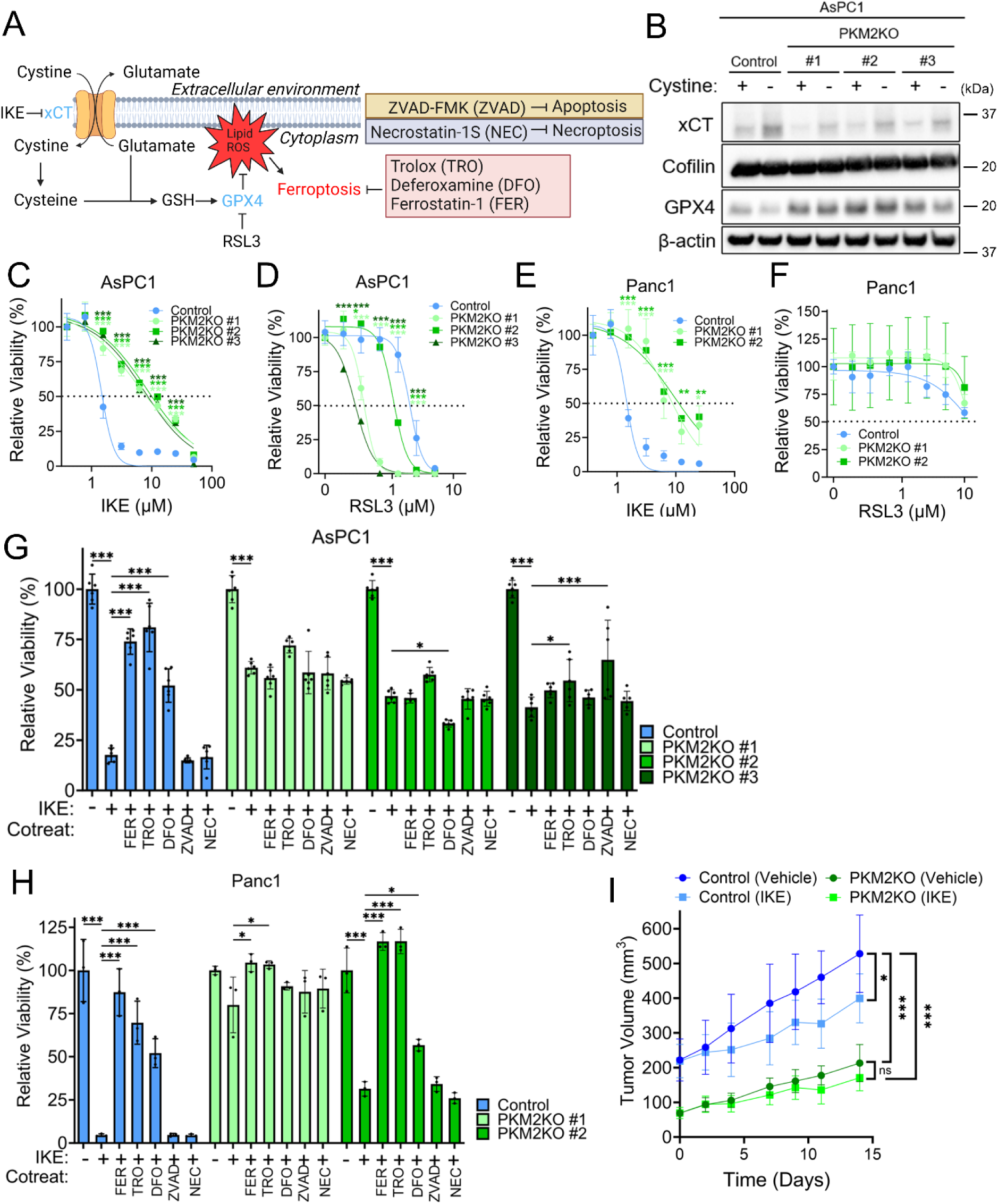
PKM2KO enhances defense against ferroptosis induction specific to cystine starvation. **A.** Schematic of the mechanism of ferroptosis, the defense proteins xCT and GPX4 (the targets of imidazole ketone erastin (IKE) and Ras selective lethal 3 (RSL3), respectively), and the ferroptosis, apoptosis, and necroptosis inhibitors used in the study. **B.** Western blot of xCT and GPX4 expression in AsPC1 control and PKM2KO cells under 50 μM (+) and 0 μM (-) cystine. **C, E.** Concentration dependent response in viability of AsPC1 (**C**) and Panc1 (**E**) control and PKM2KO clones to a range of IKE concentrations from 50-0 μM. **D, F.** Concentration dependent viability responses of AsPC1 (**D**) and Panc1 (**F**) control and PKM2KO clones to a range of RSL3 concentrations from 10-0 μM. For **C**-**F**, significance is determined two-way ANOVA. \**p*<0.05, \*\*\**p*<0.01, \*\*\**p*<0.001. Multiple hypothesis correction by the Dunnet test. **G, H.** Relative viabilities of AsPC1 (**G**) and Panc1 (**H**) control and PKM2KO cells under 50 μM cystine with 5 μM IKE (+) co-treated with 5 μM ferrostatin-1 (FER), 100 μM trolox (TRO), 100 μM deferoxamine (DFO), 50 μM Z-VAD-FMK (ZVAD), or 10 μM necrostatin-1S (NEC). Significance was assessed by two-way ANOVA. \**p*<0.05, \*\*\**p*<0.001. Multiple hypothesis correction by Tukey test. **I.** Growth of xenograft tumors produced from AsPC1 control and PKM2KO cells treated with vehicle control or 50 mg/kg IKE. Significance was assessed by two-way ANOVA at end point. \**p*<0.05, \*\*\**p*<0.001, ns = non-significant. Multiple hypothesis correction by Tukey test.

We further explored the difference in ferroptosis defense between control and PKM2KO cells by evaluating their dose response to IKE and RSL3. Fascinatingly, both AsPC1 and Panc1 PKM2KO cells demonstrate a significantly high degree of resistance to IKE compared to controls; however, these same cells have equivalent or enhanced sensitivity to RSL3 (Fig. 4C-F). We found that while the control cells treated with IKE were significantly rescued by co-treatment with ferrostatin-1, trolox, or deferoxamine, the PKM2KO clones largely show little to no response to these rescue agents (Figs. 4G-H and S4A). Additionally, there were no rescue effects from Z-VAD-FMK nor necrostatin-1S indicating that the mechanism of cell death induced by IKE is specific to ferroptosis (Fig. 4G-H). AsPC1 PKM2KO cells show decreased lipid peroxidation under IKE and equivalent lipid peroxidation under RSL3 compared to control cells (Fig. S4D). Additionally, all AsPC1 cells show strong sensitivity to RSL3 that can only be rescued by ferroptosis inhibitors, while Panc1 cells show limited sensitivity to RSL3 (Figs. S4B and E). This suggests that PK is not uniformly responsible for providing defense against all routes to ferroptosis, but rather specifically influences the metabolic reprogramming that occurs under cystine starvation and the subsequent propensity towards ferroptosis.

We next investigated whether this difference in IKE sensitivity would have the same effect on control and PKM2KO cells *in vivo*. We generated xenograft tumors in NSG mice using AsPC1 control and PKM2KO clone #1 cell lines. PKM2KO cell tumors had significantly lower tumor growth compared to the control (Fig. 4I). IKE treatment produced a significant decrease in tumor volume in the PKM2 expressing control tumors in contrast to the limited response in the PKM2KO tumors (Fig. 4I). Therefore, PKM2KO in PDAC provides resistance to cystine starvation induced cell death *in vitro* and *in vivo*.

### PKM2KO PDACs exhibit increased glutamine anaplerosis and decreased glucose metabolism under cystine starvation

PKM isoform selection is known to cause reprogramming in the overall metabolic activity within cancer cells, including both glucose and glutamine metabolism.^16–20^ To evaluate changes in metabolic pathways under low cystine conditions that may explain the survival advantage in PKM2KO cells, we performed targeted mass-spectrometry and stable isotope tracing. Using ^13^C_1,2_-glucose, we traced the flow of glucose-derived carbon through glycolysis and into the TCA cycle. The abundance of labeled hexose-phosphate is equivalent between control and PKM2KO cells under high and low cystine conditions (Fig. 5A, Fig. S5G, R). Downstream lactate production is significantly higher in control cells, which is consistent with PKM2’s role in mediating increased glucose fermentation to lactate, even in the presence of oxygen (Warburg effect).^42,43^ However, labeled lactate production significantly decreases under cystine starvation, suggesting cystine starvation may limit the Warburg effect (Figs. 5B and Fig. S5I, T). Glucose-derived carbon entry into the TCA cycle intermediates is also lower under cystine starvation, consistent with the observation that PK activity is decreased under these conditions (Figs. 5C-F and S5J-M, U-X).

**Figure 5.**
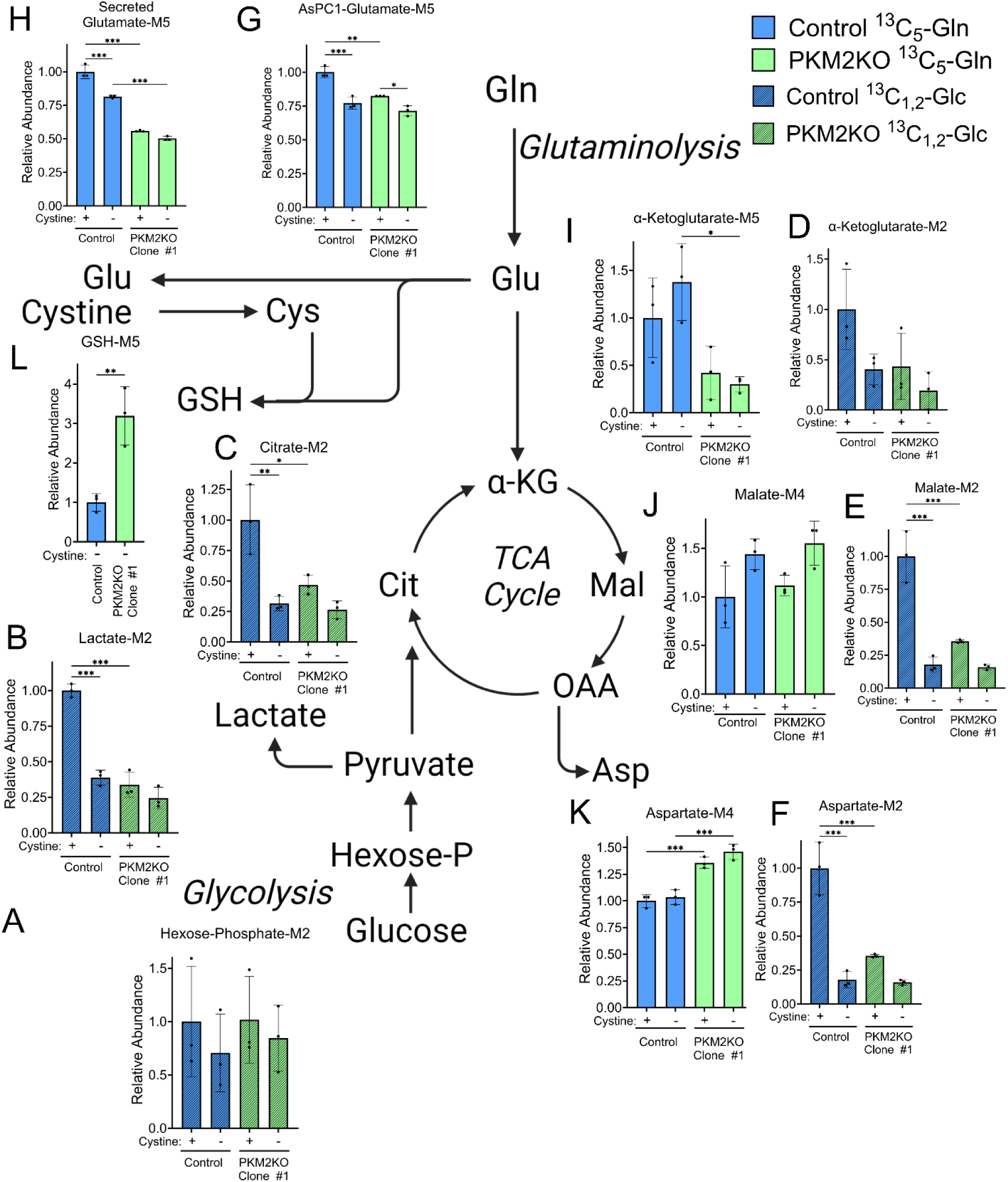
PKM2KO PDACs exhibit increased glutamine anaplerosis and decreased glucose metabolism under cystine starvation. **A-F.** Stable isotope tracing of ^13^C_1,2_-glucose under 50 μM (+) and 0 μM (-) cystine for 4 hours in AsPC1 control and PKM2KO clone #1 to produce M+2 labeled hexose-phosphate (**A**), lactate (**B**), citrate (**C**), α-ketoglutarate (**D**), malate (**E**), and aspartate (**F**). **G-K** Stable isotope tracing of ^13^C_5_-glutamine under 50 μM (+) and 0 μM (-) cystine for 24 hours in AsPC1 control and PKM2KO clone #1 to produce M+5 labeled glutamate (**G**), secreted glutamate (**H**), α-ketoglutarate (**I**), glutathione (**L**), M+4 labeled malate (**J**), and aspartate (**K**). Significance was assessed by two-way ANOVA. \**p*<0.05, \*\**p*<0.01, \*\*\**p*<0.001. Multiple hypothesis correction by Tukey test.

Given the decrease in glucose metabolism, we hypothesized that the cells were turning to alternative sources for metabolic fuel. Glutamine is an obvious candidate, as KRAS mutated pancreatic cancer cells rely on this metabolite,^44–46^ and mitochondrial glutaminolysis contributes to ferroptosis.^30^ Given this connection, we hypothesized that PKM2 expression alters glutamine metabolism and promotes ferroptosis. Thus, we next supplied the cells with uniformly (U) labeled U-^13^C_5_-glutamine and observed its utilization in downstream pathways. We first found that the breakdown of glutamine into glutamate is slightly decreased in PKM2KO cells and under cystine starvation (Figs. 5G and S5A). PKM2KO cells also secrete significantly lower glutamate than the PKM2 expressing controls (Figs. 5H and S5B). Additionally, PKM2KO cells have significantly lower production of α-ketoglutarate from glutamine while having largely equivalent amount of malate and succinate production downstream in the TCA cycle (Figs. 5I-J and S5C-F, N-Q). We also observe that PKM2KO cells have increased glutamine contribution to glutathione (Fig. 5L) and amino acid synthesis, including aspartate and proline (Fig. 5K and S8A, C). Asparagine synthesis from glutamine is significantly lower in PKM2KO cells under high cystine (Fig. S8B). However, asparagine synthesis from glutamine dramatically increases under cystine starvation, suggesting that asparagine synthesis may be involved in an important metabolic defense system against cystine starvation induced ferroptosis (Fig. S8B). Importantly, we did not observe increased glucose contributions to aspartate or alanine in the PKM2KO cells (Fig. S8D-E), suggesting that there is not a global increase in amino acid synthesis, but rather a specific utilization of glutamine for production of these amino acids. Together, these observations show that under cystine starvation, there is metabolic reprogramming away from glucose metabolism with a persistent reliance on glutamine metabolism, and that the absence of PKM2 influences how glucose and glutamine are metabolized.

We additionally used these metabolic tracer studies to address other hypotheses that might explain the increased viability of PKM2KO cells under cystine starvation. Decreased PK activity has been linked to increased flux through the pentose phosphate pathway (PPP), an important pathway for generating NADPH by glucose-6-phospate dehydrogenase.^33^ Since NADPH is a critical reducing agent for reducing glutathione (GSH) and defending against lipid peroxidation, this raises the possibility that utilization of the PPP is more active in the PKM2KO cells under low cystine stress. To address this, we used the ^13^C_1,2_-glucose tracer to take advantage of the fact that flux through the PPP results in the loss of the first carbon. Subsequent re-entry into glycolysis would result in M1 labeling, while flux directly through glycolysis will produce M2 labeling downstream. We observe that M1 labeling in ribose-5-phosphate and ribulose-5-phosphate is actually decreased under cystine starvation, though the result was not significant (Fig. S6A-D). Further, there were no significant differences between control and PKM2KO cells. We did observe significantly more M1 isotopologue hexose-phosphate in one AsPC1 PKM2KO clone, but no consistent or significant trend in the other cells (Fig. S6E-F). This suggests that differential utilization of the PPP is likely not a major driver for explaining the difference in defense against ferroptosis between PKM2KO cells and PKM2 expressing cells.

Glutathione (GSH) is a critical antioxidant defense molecule, synthesized from cysteine, glycine, and glutamate.^47^ Given the difference in glutamine metabolism between control and PKM2KO cells under cystine starvation, we hypothesized that the PKM2KO cells would have higher GSH synthesis. We used U-^13^C_5_-glutamine to trace the flow of glutamine derived carbon into GSH, to produce the M5 isotopologue of GSH. As expected, GSH pool size plummets under cystine starvation, likely due to rapid consumption for quenching lipid peroxides and the lack of new cysteine to synthesize more (Fig. S7A, C). Although the pool of GSH is extremely low, we do observe significantly increased M5 GSH in two of the AsPC1 PKM2KO clones under cystine starvation, but this effect was not seen in the Panc1 cells (Figs. 5L and S7B, D). To further explore potential differences in GSH synthesis, we co-treated the control and PKM2KO cells with buthionine sulfoximine (BSO), an inhibitor of glutathione synthesis that does not induce ferroptosis.^22,48^ However, we observe a minor decrease in viability with BSO treatment that can be restored by co-treatment with ferrostatin-1 in AsPC1 cells but not Panc1 cells (Fig. S7E-G). Under cystine starvation, BSO treatment does not alter viability of either control or PKM2KO cells, indicating that inhibition of GSH synthesis cannot further alter ferroptosis when environmental cystine is removed. Together, these results suggest that there are some minor differences in GSH synthetic capabilities between control and PKM2KO cells, but the ability to synthesize GSH is likely not a major contributor to the difference in ability to defend against ferroptosis.

Lastly, we hypothesized that the increased production of amino acids from glutamine in the PKM2KO cells was providing the cells with additional resources to maintain viability under cystine starvation. We removed environmental cystine and supplemented the cells with supraphysiologic concentrations of aspartate, asparagine, glutamate, and proline individually. However, this produced no change in viability compared to cystine starvation alone and did not alter the difference in viability between control and PKM2KO cells (Fig. S8F). This suggests that the ability of the PKM2KO cells to increase production of amino acids from glutamine is not essential for improving cellular defense against ferroptosis. However, it should be noted that directly supplying the end products (amino acids) is not equivalent to synthesizing them, as the reactions to produce these products also affect redox cofactors and stoichiometries of the reactants. Altogether, this data suggests a complex utilization of glutamine metabolism by PKM2KO cells to support defense against cystine starvation induced ferroptosis.

### Glutamine is required for PKM2KO PDAC defense against cystine starvation induced ferroptosis

Given the apparent importance of glutamine metabolism in ferroptosis, we investigated the role of environmental glutamine in PKM2KO mediated defense against ferroptosis. Glutamine metabolism is an important component of ferroptosis.^30^ Given the difference in glutamine metabolism in the PKM2KO cells, we tested whether environmental glutamine availability would influence cell survival. The difference in viability under cystine starvation between PKM2KO and control cells is present with glutamine levels as low as 250 μM, but complete removal of glutamine significantly decreases viability of the PKM2KO cells and restores viability to control cells (Fig. S9A). This is consistent with our observation that PKM2KO cells have elevated glutamine metabolism, and therefore increased dependence, on glutamine. We found that complete removal of glutamine significantly enhances the viability of the control cells under cystine starvation. In contrast, the PKM2KO cells have reduced viability when glutamine is removed under cystine starvation induced by IKE treatment (Fig. 6A-B). Additionally, we observed the same effect when the cells are grown under 0 μM cystine starvation mediated ferroptosis (Fig. 6C). Glutamine metabolism can also be inhibited by the glutaminase inhibitor CB-839.^49,50^ Treatment with CB-839 significantly decreases the viability of PKM2KO cells, but has a more muted effect on the control cells (Fig. S9B). In contrast, under cystine starvation, CB-839 has no effect on viability in the PKM2KO cells, while restoring viability in the control cells (Fig. S9B). We further observe that removal of glutamine eliminates the increased lipid peroxidation observed in control cells under 0 μM cystine conditions and that glutamine removal causes no significant increase in lipid ROS in either control or PKM2KO cells compared to replete media conditions (Fig. 6D). We next evaluated whether simultaneous glutamine and cystine starvation would alter expression of the ferroptosis defense proteins, xCT and GPX4. Removal of cystine causes an increase in xCT expression and decrease in GPX4 expression in the AsPC1 and Panc1 control cells, but only modest changes in the PKM2KO cells (Fig. 6E-F). When both environmental glutamine and cystine are removed, we no longer observe the increase in xCT or decrease in GPX4, indicating that either the stressful stimuli promoting their expression is absent or the presence of glutamine is required to promote expression (Fig. 6E-F). Together, these results support the conclusion that altered glutamine metabolism in PKM2KO cells is in part responsible for the differential ferroptosis response under cystine starvation conditions.

**Figure 6.**
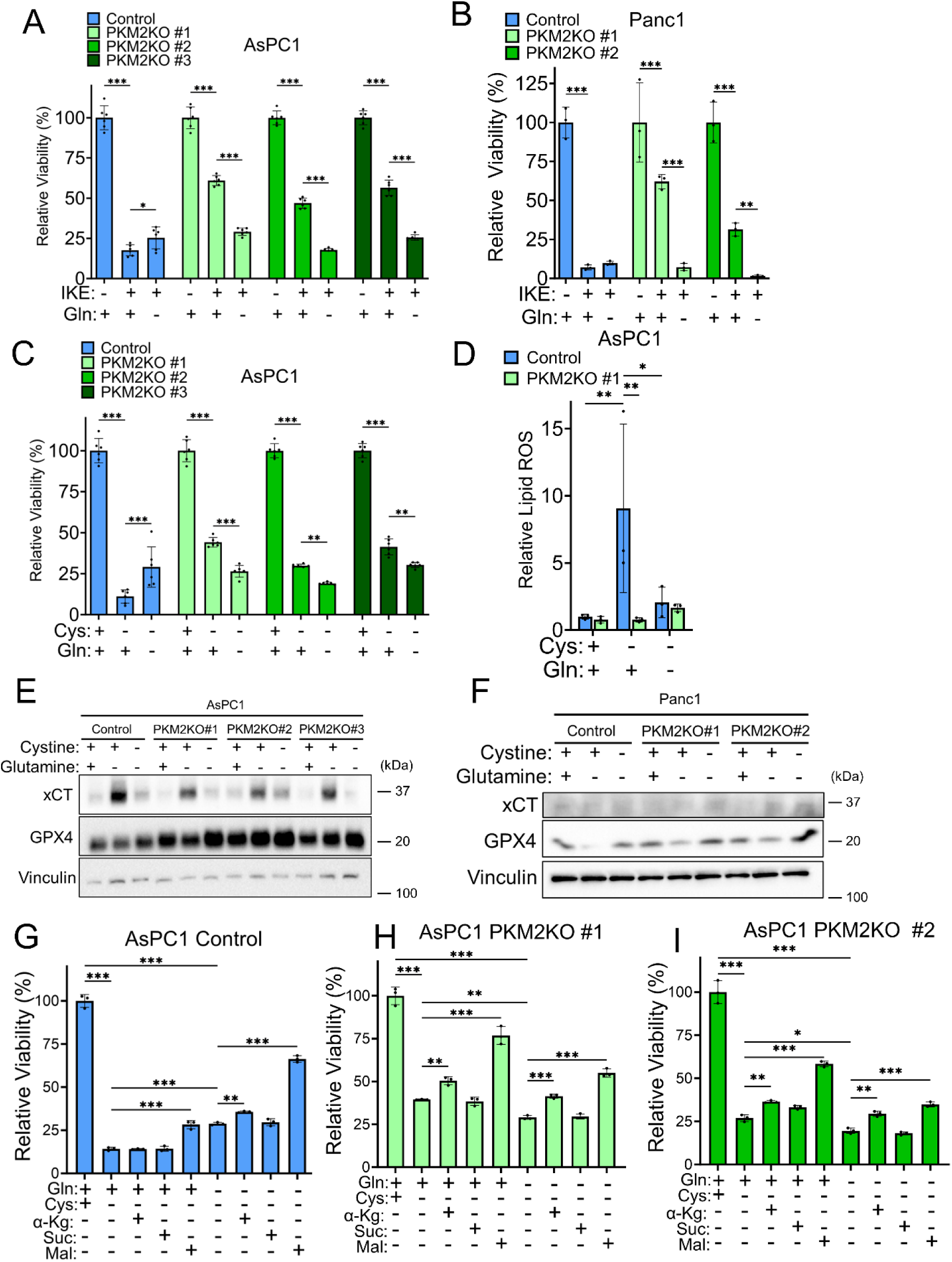
Glutamine is required for PKM2KO PDAC defense against cystine starvation induced ferroptosis. **A, B.** Relative viabilities of AspC1 (**A**) and Panc1 (**B**) control and PKM2KO cells under 50 μM cystine with (+) or without (-) 1 mM glutamine treated with 5 μM IKE. **C.** Relative viabilities of AsPC1 control and PKM2KO cells under 50 μM (+) and 0 μM (-) cystine with (+) or without (-) 1 mM glutamine. **D.** Relative lipid peroxidation in AsPC1 Control and PKM2KO clone #1 under 50 μM (+) or 0 μM (-) cystine with (+) or without (-) 1 mM glutamine. For **A**-**D**, significance was assessed by two-way ANOVA. \**p*<0.05, \*\**p*<0.01, \*\*\**p*<0.001. Multiple hypothesis correction by Tukey test. **E, F.** Western blot of xCT and GPX4 expression in AsPC1 (**A**) and Panc1 (**F**) cells under 50 μM (+) and 0 μM (-) cystine with (+) or without (-) 1 mM glutamine. **G-I.** Relative viabilities of AsPC1 control and PKM2KO clones under 50 μM (+) or 0 μM (-) cystine with (+) or without (-) 1 mM glutamine supplemented with either 8 mM dimethyl-α-ketoglutarate (αKG), 8 mM dimethyl-succinate (Suc), or 32 mM dimethyl-malate (Mal). Significance was assessed by one-way ANOVA. \**p*<0.05, \*\**p*<0.01, \*\*\**p*<0.001. Multiple hypothesis correction by Sidak test.

Glutamine anaplerosis is an important metabolic process for filling the resources required for the TCA cycle to function in the mitochondria of cells in the low nutrient conditions of the PDAC tumor microenvironment^18,21^ and is influenced by PK enzyme activity.^16,51^ To address whether metabolites downstream of glutaminolysis would influence ferroptosis, we grew the AsPC1 and Panc1 PKM2KO cells in the absence of cystine and co-treated the cells with cell permeable dimethyl α-ketoglutarate (DM-αkg), dimethyl succinate (DM-Suc), and dimethyl malate (DM-Mal) at concentrations shown to increase ferroptosis.^30^ In the AsPC1 cells, the addition of DM-αkg or DM-Suc did not change viability. Fascinatingly, the addition of malate significantly restored viability to these cells (Fig. 6G-I and S9C). Even more surprisingly, when environmental glutamine is removed, we observe the same trend where DM-αkg and DM-Suc do not affect viability, and treatment with DM-Mal further increases viability (Figs. 6G-I and S9C). The AsPC1 PKM2KO cells have worse viability when environmental glutamine is removed, yet the addition of DM-αkg or DM-Mal, but not DM-Suc, rescues viability loss caused by glutamine removal (Fig. 6G-I and S9C). Treatment with DM-Mal also restores viability when glutamine metabolism is blocked by CB-839 (Fig. S9B). While this effect is prominent in the AsPC1 cells, we do not observe the same effect in Panc1. The viability of Panc1 cells under cystine starvation is restored by DM-αkg, but further decreased when DM-Suc and DM-Mal are supplemented (Fig. S9D-F). To confirm that this is not an artifact of higher concentration of malate, we also find that using a lower concentration of DM-Mal (8 mM) is sufficient to cause the same changes in the AsPC1 and Panc1 PKM2KO cells (Fig. S9G-H). Supplementation with DM-αkg and DM-Suc produce little change under cystine and glutamine replete conditions, but DM-Mal supplementation consistently increases viability (Fig. S9I-O). Collectively, our data presents strong evidence that PK plays a key role in altering ferroptosis by reprogramming glutamine metabolism and suggests that under low PK activity, cells utilize enhanced glutamine metabolism under low cystine conditions for survival.

### Malic enzyme enables survival of PKM2KO PDAC under cystine starvation

We next aimed to identify the metabolic pathway that the PKM2KO cells rely on to alter glutamine metabolism and promote defense against ferroptosis. KRAS mutant PDAC upregulates several metabolic enzymes, including glutamate oxaloacetate transaminase 1 (GOT1), glutamate oxaloacetate transaminase 2 (GOT2), and malic enzyme 1 (ME1), to alter glutamine metabolism and promote defense against ferroptosis.^46^ Additionally, inhibition of GOT1 promotes ferroptosis in pancreatic cancer.^52^ The fact that malate promotes increased survival of cystine starvation and that our U-^13^C_5_-glutamine labeling data demonstrates high levels of glutamine flux into malate and aspartate led us to hypothesize that malic enzyme and the malate aspartate shuttle are important for providing the metabolic advantage in PKM2KO cells under cystine starvation. To address this possibility, we evaluated the expression of ME1 in the AsPC1 and Panc1 PKM2KO. Surprisingly, the PKM2KO clones demonstrate elevated ME1 expression compared to their respective controls; however, expression was minimally impacted by the presence or absence of cystine (Fig. 7A-B). We next attempted to block ME1 using malic enzyme inhibitor (ME1i) to test whether this would influence ferroptosis. We found that inhibition of ME1 significantly decreases viability of control and PKM2KO cells under cystine starvation (Figs. 7C-G and S10A-B) Co-treatment with ferrostatin-1 was able to significantly restore viability in the Panc1 cells (Fig. 7C-E). In AsPC1 cells, only N-acetylcysteine (NAC), an antioxidant, was able to rescue this effect consistently (Figs. 7F-G and S10A-B). This indicates that the decrease in viability induced by cystine starvation and ME1 inhibition in AsPC1 cells is likely specific to oxidative stress but may not be entirely explained by ferroptosis and speaks to the metabolic heterogeneity that has been observed in pancreatic cancer.^2^ We further demonstrate that supplementation with malate in addition to ME1 inhibition ablates the viability advantage that malate supplementation alone provides in AsPC1 cells (Figs. 7H-I and S10C-G). Because ME1 is important for generating the antioxidant NADPH, we next evaluated NADPH levels in the PKM2KO cells. We observe that NADPH levels are consistently higher in the PKM2KO cells under 0 μM cystine, consistent with increased defense against ferroptosis (Fig. 7J-K). Based on these results, we have developed a model in which PK influences the expression of ME1 and that under low PKM2 activity, PDAC cells increase glutamine metabolism and utilization of ME1 to promote NADPH production and defense against ferroptosis (Fig. 7L). Furthermore, activation of PKM2 increases glycolysis and de-emphasizes utilization of glutamine for NAPDH generation and promotes metabolic conditions more susceptible to ferroptosis (Fig. 7M).

**Figure 7.**
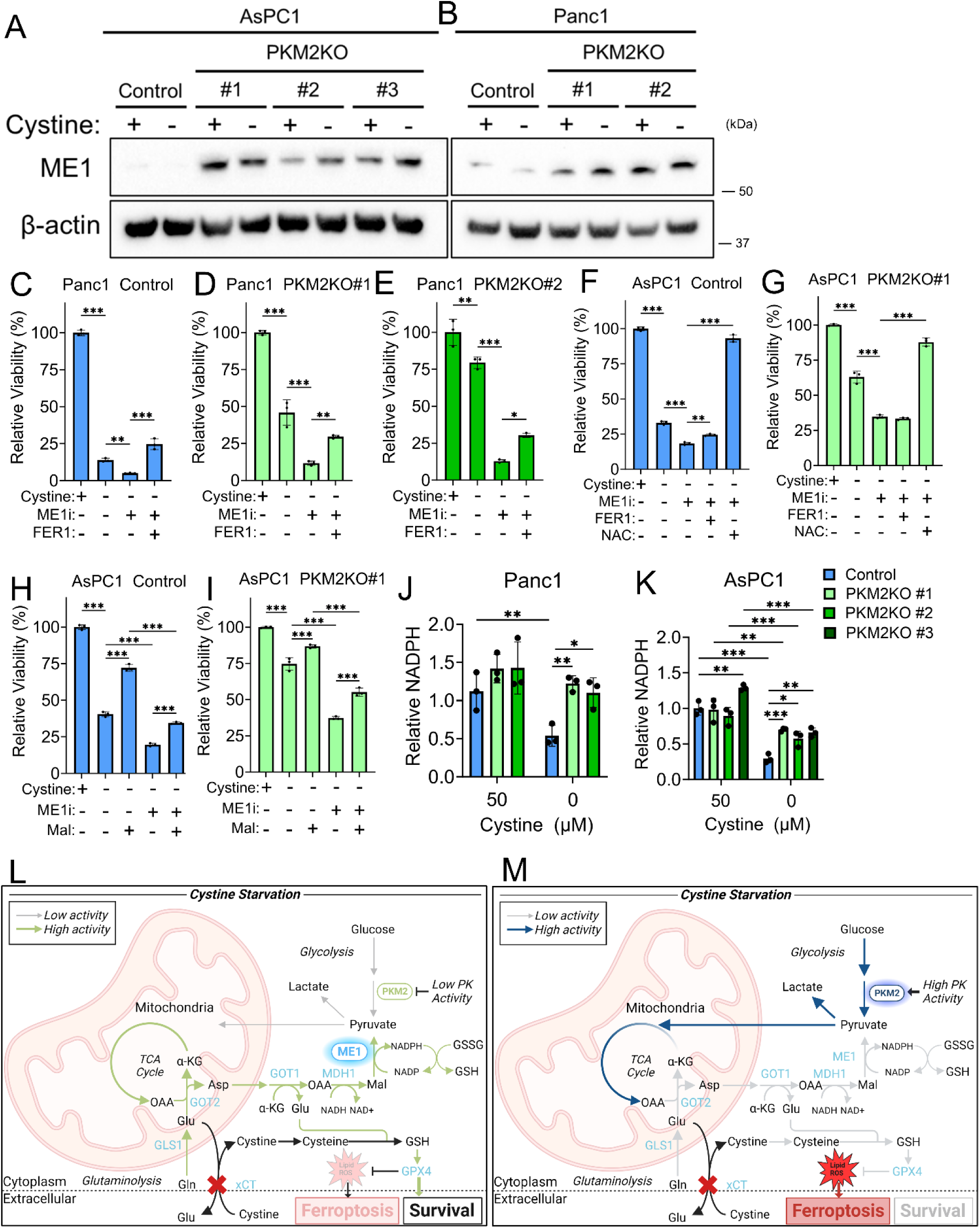
Malic enzyme enables survival of PKM2KO PDAC under cystine starvation. **A, B.** Western blot of malic enzyme 1 (ME1) expression in AsPC1 (**A**) and Panc1 (**B**) control cells and PKM2KO clones under 50 μM (+) or 0 μM (-) cystine conditions. **C-G**. Relative viabilities of Panc1 control cells (**C**), Panc1 PKM2KO #1 (**D**), Panc1 PKM2KO #2 (**E**), AsPC1 control cells (**F**), and AsPC1 PKM2KO clone #1 (**G**) under 0 μM cystine treated with (+) or without (-) 50 μM malic enzyme 1 inhibitor (ME1i) and co-treated with either 5 μM ferrostatin-1 (FER) or 1 mM N-acetylcysteine (NAC). **H, I.** Relative viabilities of AsPC1 control (**H**) and PKM2KO clone #1 (**I**) under 50 μM (+) or 0 μM (-) cystine with (+) or without (-) 50 μM ME1i and 32 mM dimethyl-malate (Mal) supplement. For **C**-**I**, significance was assessed by one-way ANOVA. \**p*<0.05, \*\**p*<0.01, \*\*\**p*<0.001. Multiple hypothesis correction by Sidak test. **J, K.** Relative NADPH abundance in AsPC1 (**J**) and Panc1 (**K**) control and PKM2KO clones under 50 or 0 μM cystine. For **J**-**K**, significance was assessed by two-way ANOVA. \**p*<0.05, \*\**p*<0.01, \*\*\**p*<0.001. Multiple hypothesis correction by Tukey test. **L-M.** Proposed model on how PK reprograms metabolism to influence cystine starvation induced ferroptosis under low PK activity (**L**) and high PK activity (**M**).

### The combination of pyruvate activation and cystine starvation is an efficacious treatment for PDAC *in vivo*

Finally, we tested the impact of increasing pyruvate kinase activity and inhibiting xCT on PDAC growth *in vivo* using NOD scid gamma (NSG) mice injected with Panc1 WT cells. Mice were divided into 4 treatment groups to receive either 50 mg/kg IKE, 30 mg/kg TEPP-46, combination of 50 mg/kg IKE and 30 mg/kg TEPP-46, or vehicle control (Fig. 8A). After 2 weeks of daily intraperitoneal injections, tumor volume was significantly lower in the TEPP-46 and combination treatment groups compared to the vehicle control group (Fig. 8B). Tumor weight was also significantly lower in the combination treatment group compared to the control and TEPP-46 treatment groups, with a trend towards decreased weight compared to IKE treatment group (Fig. 8C-D). No significant differences in body weight were observed between each of the groups over the duration of the treatment. This provides proof-of-concept that activating pyruvate kinase and inducing cystine starvation is a viable, novel treatment strategy for PDAC.

**Figure 8.**
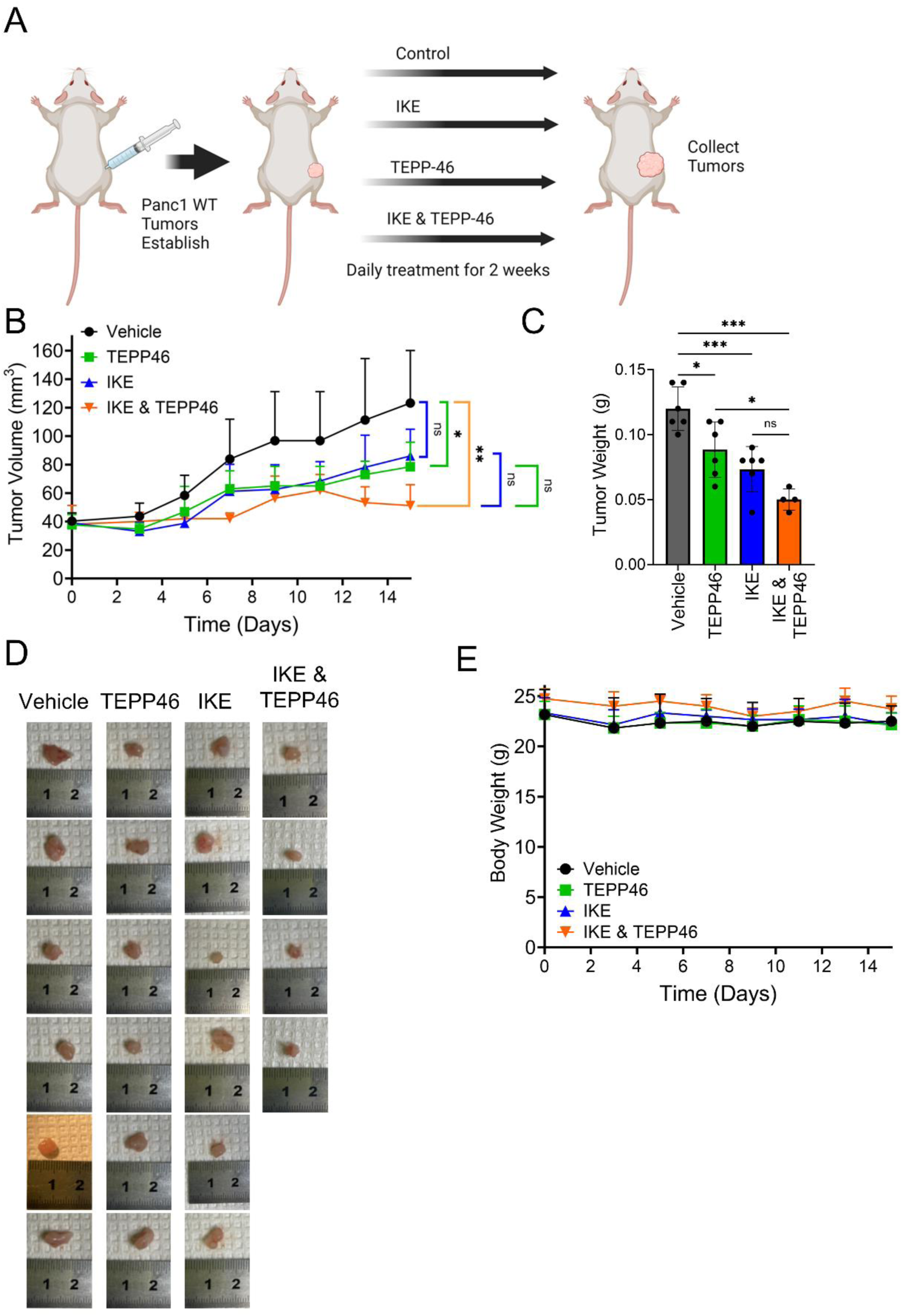
The combination of pyruvate activation and cystine starvation is an efficacious treatment for PDAC *in vivo*. **A**. Treatment schematic for xenograft tumors formed from Panc1 WT cells, treated daily for 2 weeks with vehicle control, IKE, TEPP-46, or IKE and TEPP-46 combined. N=6 for each treatment group, except IKE and TEPP-46 combination with N=4. **B.** Tumor volume of xenograft tumors for each treatment group over time. **C**. Tumor weight at end point for each treatment group. For **B**-**C**, significance was assessed by one-way ANOVA at end point. \**p*<0.05, \*\**p*<0.01, ns = non-significant. Multiple hypothesis correction by Sidak test. **D**. Images of tumors for each treatment group at end point. **E.** Body weight of treated mice throughout the treatment course.

## Discussion

Pyruvate kinase plays a complex role in the metabolic reprogramming of pancreatic cancer cells. Here, we demonstrate for the first time that PKM2KO provides a survival advantage against cystine starvation induced ferroptosis. We further show that this survival advantage is mediated by decreased PK activity that leads to reprogrammed glutamine metabolism and activation of ME1 to produce NADPH. Ferroptosis is inherently a product of aberrant metabolic conditions and represents an exciting new avenue of targeted metabolic therapy.^2,53^ PDAC tumor cells survive under low nutrient conditions and are particularly vulnerable to cystine starvation induced ferroptosis, which has been recently demonstrated *in vivo*.^22^ Our study provides evidence that decreasing PK activity is a potential resistance mechanism against cystine starvation induced ferroptosis. While increased PK activity sensitizes cells to oxidative stress, its activity has not yet been connected to ferroptosis.^33,54^ Here, we demonstrate for the first time that activation of PKM2 synergistically enhances cystine starvation induced ferroptosis and suggests a novel strategy for inhibiting survival of pancreatic cancer cells. It has recently been proposed that there is no single unified method for ferroptosis; instead, it exists as a flexible penumbra of regulated defenses.^55^ Consistently, we find that decreased PK activity does not universally protect against all ferroptosis, but specifically against cystine starvation induced ferroptosis, as inhibition of GPX4 leads to ferroptosis even with low PK activity.

We demonstrate that the combination of cystine starvation and activation of PKM2 is a novel and efficacious therapy for treating PDAC *in vitro* and *in vivo*. This newly discovered metabolic vulnerability in PDAC represents a promising opportunity for therapeutic intervention in pancreatic cancer. Further research will identify and optimize potent and selective drugs for treatment of PDAC. Interestingly, IKE treatment alone had only a modest effect on PDAC tumor growth in contrast to xenograft tumors from other cancer types.^29,56^ Our findings also raise the possibility that cancer cells with low PKM2 expression or low PK activity may provide a pool of persistent cells within a tumor that resist cystine starvation. Thus, the combination of increasing pyruvate kinase activity and inducing cystine starvation presents exciting opportunities for targeting persistent cancer cells.

Our work provides further clarity on the role of glutamine metabolism in ferroptosis, which has been somewhat controversial, in PDAC. In other cancer types, removal of glutamine suppressed ferroptosis induced by cystine starvation.^27,30,57^ On the other hand, recent observation in pancreatic and hepatocellular cancer cell models show that the removal of glutamine can possibly cause ferroptosis.^58,59^ Here, we report that the removal of glutamine eliminates lipid peroxide accumulation and prevents ferroptosis only in PKM2 expressing PDAC cells under cystine starvation. In contrast, removal of glutamine or inhibition of glutaminase decreases survival of PKM2KO cells under cystine starvation. To further improve our understanding of glutamine metabolism in ferroptosis in PDAC, we employed ^13^C-glutamine tracing to evaluate glutaminolysis under cystine starvation conditions. The PKM2KO cells show resistance to ferroptosis and simultaneously active glutaminolysis through producing TCA cycle intermediates and deriving amino acids from glutamine. This suggests there are metabolic states in which glutamine can be used in either a pro- or anti-ferroptotic manner. We propose that PK plays a key role in coordinating glucose and glutamine metabolism as a potential adaptation response to cystine starvation and resistance to ferroptosis.

Investigating metabolic pathways known to influence antioxidant defense revealed that neither the oxidative pentose phosphate pathway nor glutathione synthesis is the dominant mechanism by which PKM2KO cells defend against ferroptosis. Rather, malic enzyme and the malate aspartate shuttle are important for providing the metabolic advantage in PKM2KO cells under cystine starvation, as evidenced by malate supplementation promoting survival of cystine starvation, increased U-^13^C_5_-glutamine flux into malate and aspartate, and elevated ME1 expression in PKM2KO cells. KRAS mutated cancer cells, such as PDAC cells, decouple glucose and glutamine metabolism and upregulate ME1 in addition to GOT1 and GOT2 to facilitate the breakdown of glutamine for anaplerosis and production of NADPH.^45,46,52^ ME1 has also been identified as a ferroptosis defense protein in hepatic ischemia/reperfusion injury and synovial sarcoma.^60,61^ Our work establishes a novel connection between PK and ME1 as well as their roles in ferroptosis in pancreatic cancer. Additionally, our study shows that inhibition of ME1 provides a route to promote ferroptosis and circumvent PDAC metabolic defense strategies for surviving low cystine conditions.

It has been established that the PDAC tumor microenvironment is deficient in many nutrients, including cystine.^38^ The mechanism driving this low cystine environment is currently unclear, but possible explanations include changes in tumor cell genetics or stromal cell composition within the tumor.^38^ In our study, we have intentionally used glucose, glutamine, and cystine concentrations reflective of the tumor microenvironment. The low cystine environment has also been shown to influence central carbon metabolism. Specifically, lung cancer cells grown under low cystine conditions have decreased glutamine metabolism and conversely supraphysiologic concentrations of cystine drive enhanced glutaminolysis.^62^ The requirement of high cystine to drive glutamine metabolism may explain the lack of robust results in targeting glutamine metabolism in clinical trials.^63,64^ The expression of the cystine/glutamate antiporter, xCT, also enhances dependency on glucose as increased glutamate is secreted rather than used in downstream metabolic pathways.^65^ Further, high expression levels of xCT increases sensitivity to oxidative stress and alters glutamine metabolism,^66,67^ consistent with our observation that PKM2KO cells have decreased xCT expression under cystine starvation.

Upstream stimulatory factor 2 inhibits erastin induced ferroptosis through transcriptional regulation of PKM2 in pancreatic cancer.^68^ This study showed that knockdown of PKM2 enhances sensitivity to erastin induced ferroptosis. This difference from our results likely comes from our complete deletion of PKM2, which puts the cells in a fundamentally different metabolic state than decreasing its expression by knockdown. As the previous work did not assess the role of PK activity or utilize metabolic tracing, our work expands on these findings and provides a more robust picture of the role of PK as it relates to ferroptosis in pancreatic cancer.

The precise mechanism by which PKM2 activity changes expression of ME1, xCT, and GPX4 has yet to be determined. Decreased PK activity leads to decreased pyruvate production. As malic enzyme uses malate to produce pyruvate, upregulation of ME1 may serve as a compensatory route for maintaining adequate pyruvate pools. PKM2 mediated reprogramming of metabolism has known effects on nutrient sensing, accumulation of biosynthetic intermediates, and production of antioxidants.^16–19,33,51,69–71^ These changes themselves likely have downstream signaling effects leading to complex changes in metabolic machinery and defense systems against ferroptosis. Additionally, dimeric PKM2 translocates into the nucleus and influences transcription of several genes implicated in cancer progression including Cyclin D1, C-Myc, HIF-1α, and β-catenin.^10,72,73^ Some of these factors are known to impact ferroptosis; for example, β-catenin is involved in GPX4 synthesis.^74^ PKM2 can also inhibit p53, which is known to decrease xCT expression, providing a potential connection between PKM2 and xCT.^75–78^ HIF-1α promotes increased iron uptake and availability suggesting potential connections between iron metabolism and PKM2 which would likely influence ferroptosis.^79–81^ Additionally, our finding that low PK activity protects against cystine starvation induced ferroptosis suggests that cells with low PKM2 or PKM1 expression may represent a sub-population of cells able to persist under stressful low nutrient conditions of the tumor microenvironment and allow for tumor survival.^11^ Therefore, the role of PKM2 in ferroptosis is multifaceted and worthy of further investigation.

In conclusion, our work demonstrates that decreasing PK activity protects PDACs from cystine starvation induced ferroptosis. Additionally, our data show that cancer cells with low PK activity represent a potential pool of resilient cells able to tolerate low nutrient stress and enable tumor persistence. Therefore, this work reveals critical mechanisms by which PDAC cells reprogram their metabolic pathways to support survival when environmental cystine is scarce. Targeted PKM2 activation in combination with cystine metabolism inhibition is a novel strategy against PDAC tumors and represents a promising new path to improve outcomes for pancreatic cancer patients.

### Limitations of This Study

We have limited our evaluation of the role of PK as a metabolic enzyme. However, PKM2 has additional moonlighting functions as a transcription factor, and we intend to investigate this in future experiments and identify a direct connection between PKM2 and the observed gene expression changes described here. Pancreatic cancer has a high degree of metabolic and genetic heterogeneity.^2,82–88^ Indeed, we observe metabolic differences between the AsPC1 and Panc1 cells. Specifically, supplementation with exogenous malate under cystine starvation can restore viability to the AsPC1 control and PKM2KO cells by providing fuel for malic enzyme to produce NADPH. However, malate supplementation worsens survival of Panc1 control and PKM2KO cells as observed in other cancer types.^30^ This suggests that further work is still needed to understand the metabolic heterogeneity influencing ferroptosis in PDAC and other cancer types. *In vivo* studies were conducted in heterotopic xenograft models in immunocompromised mice. Future studies will include orthotopic models in immunocompetent mice and optimize drug selection, doses, pharmacokinetics, and treatment regimen.

## Methods and Materials

### Cell culture

The human pancreatic cancer cell lines, AsPC1, Panc1, BxPC3, and MiaPaCa2 were acquired as a gift from Dr. Nouri Neamati at University of Michigan. Cells were routinely cultured in Dulbecco’s modified eagle medium (DMEM) (MT10013CV, Fisher Scientific,) with sodium pyruvate, supplemented with 10% fetal bovine serum (FBS) (13206C, Sigma-Aldrich), 1% penicillin and streptomycin (P/S) (15140122, Fisher Scientific), 5 μg/mL plasmocin (ant-mpp, Invivogen), and cultured in a humidified incubator with 5% CO_2_ at 37 °C. During experimental procedures cells were incubated in a humidified incubator with 5% CO_2_ at 37 °C in DMEM without glucose, glutamine, pyruvate, or cystine (D9815, US Biological) supplemented with 5 mM glucose, 1 mM pyruvate, and varying amounts of glutamine or cystine as indicated, 10% dialyzed FBS (F0392, Sigma-Aldrich), and 1% P/S. Cells were routinely tested for mycoplasma detection with the kit (rep-mysnc-100, Invivogen).

### Gene knockout by lentiviral CRISPR/Cas9 gene editing

CRISPR/Cas9-mediated genome editing was used to achieve PKM2 knockout with lentivirus-mediated gene expression.^89^ Guide RNAs targeting exon 10 of the PKM gene (region determinative of PKM2 expression) were designed by CRISPR DESIGN (http://crispr.mit.edu.proxy1.cl.msu.edu/) and set just before the protospacer adjacent motif (PAM), a DNA sequence immediately following the Cas9-targeted DNA sequence. A lentiviral vector was used expressing one single guide RNA, human caspase 9, and an antibiotic selection marker. The sgRNA sequence for PKM2 knockout plasmid vector is 5′- GTTCTTCAAACAGCTTGCGG-3′ along with puromycin resistance. The sequence for scramble control plasmid with blasticidin selection maker is 5′-GCACTACCAGAGCTAACTCA -3′. CRISPR gene editing plasmid vectors with gRNA and Cas9 co-expression were acquired from VectorBuilder. The VSVG plasmid was a gift from Bob Weinberg (Addgene plasmid # 8454; http://n2t.net/addgene:8454; RRID: Addgene 8454). The psPAX2 plasmid was a gift from Didier Trono (Addgene plasmid # 12260; http://n2t.net/addgene:12260; RRID: Addgene 12260). Vectors were amplified by transforming Stbl3 bacterial cells grown in LB broth under 100 μg/mL ampicillin for antibiotic selection. Plasmids were harvested by Midi-prep (12243, Qiagen). To produce lentivirus, HEK293T cells were seeded in 10-cm plates containing OptiMEM (110580221, Fisher Scientific) with 4% FBS. When the HEK293T cells reached ∼50% confluency, they were transfected using Lipofectamine 3000 (L3000001, Fisher Scientific) according to the manufacturer’s instructions with 10.0 μg lentivirus plasmids, 0.5 μg VSVG, and 5.0 μg psPAX2 plasmids. After 24 hours, fresh DMEM with 15% FBS and 1% P/S was added, and cells were grown for another 48 hours to generate virus. For transduction with lentivirus, the AsPC1 and Panc1 cells (1 × 10^5^ cells) were seeded in 10-cm plates. The supernatant of transfected HEK293T was collected and passed through 0.45 micron PVDF syringe filter. Five mL of the viral supernatant and 5 ml of fresh media were added to recipient AsPC1 and Panc1 cell plates with polybrene (TR1003G, Fisher Scientific) at final concentration of 4 μg/ml. The cells were cultured for 24 hours followed by adding fresh DMEM medium supplemented with 10% FBS and treated for 12 days with 2 μg/mL puromycin (A1113803, Fisher Scientific) for selection. The selected cells were then expanded and analyzed for successful gene knockout by sequencing and western blot analysis. Genomic DNA was extracted using DNeasy Blood and Tissue Kit (69506, Qiagen) to check for successful gene editing. PCR primers used to amplify the targeted region around exon 10 of PKM consisted of a forward primer (5’-GCACTTGGTGAAGGACTGGT-3’) and reverse primer (5’-AATGGACTGCTCCCAGGAC-3’). A nested primer (5’-GTGACTCTTCCCCTCCCTCT- 3’) was used for sequencing. Sequencing was completed by ACTG. Individual clones were selected by diluting the population to single cells plated in a 96-well plate and then expanding the population. PKM2 deletion for isolated clones was determined by western blot.

### PKM1 and PKM2 overexpression

Overexpression of PKM1 and PKM2 was achieved using a lentiviral gene expression purchased from VectorBuilder. The vector contained the blasticidin resistance gene and either the PKM1 or PKM2 protein coding sequence under the EF1A promoter. Lentivirus was produced and used to transduce target cells in an identical manner as described in the previous section. Transduced cells were selected using 10 μg/ml blasticidin (A1113903, Fisher Scientific). The selected cells were then expanded and analyzed for successful gene knockout by western blot analysis.

### Protein extraction and western blot analysis

Cells were plated in 10 cm plates and grown until ∼60-70% confluent. Plates were washed twice with PBS w/o calcium or magnesium (D8537, Sigma-Aldrich) and the media was replaced with experimental media. Panc1 cells were incubated for 12 hours, AspC1 were incubated for 24 hours as described above. Cell lysis for protein extract was completed using cell lysis buffer (9803, Cell signaling). Western blot analysis was carried out using standard protocols. Briefly, protein samples were diluted to 1 μg/mL, reduced with 6x Laemmli loading dye, and boiled for 5 minutes. 30 μg of protein was loaded to precast Bolt 10- or 17-well 4-12% polyacrylamide, Bis-Tris, 1.0 mm gels (NW04122BOX, Fisher Scientific,). Gel electrophoresis was carried out in MES running buffer at 200V for 22 minutes. Protein was transferred to nitrocellulose membranes, which were then stained with Ponceau stain to confirm total protein equivalence between samples, and were then cut for appropriate targets of interest. The following dilutions of primary commercial antibodies in 5% BSA in tris buffered saline with 0.1% TWEEN20 (TBST) were used as probes: 1:1000 dilution of anti-PKM1 (7067S, Cell Signaling Technology), 1:1000 dilution of anti-PKM2 (4053S, Cell Signaling Technology), 1:1000 dilution of anti-GPX4 (52455, Cell Signaling Technology), 1:500 dilution of anti xCT (D2M7A, Cell Signaling Technology), 1:1000 dilution of anti-β-actin (49070S, Cell Signaling Technology), and 1:1000 dilution of anti-vinculin (13901, Cell Signaling Technology). Primary antibodies were diluted in 5% BSA in TBST and incubated overnight at 4 °C. HRP-linked secondary anti-rabbit antibodies (7074S, Cell Signaling Technology) were diluted in 5% non-fat milk at a dilution of 1:5000 and incubated at room temperature for 1 h. Clarity Western ECL Substrate Kit (1705060, BioRad) was used for detection. Images were captured on the BioRad ChemiDoc XRS+ imager and Image Lab 5.2.1 was used for image processing.

### Cell viability and proliferation analysis

Cells were plated on 96-well plates at 2000 cells/well for Panc1 and 4000 cells/well for all other cell lines under standard conditions described above. Cells were allowed to seed overnight (approximately 18 hours). Media was aspirated and cells were washed twice with phosphate buffered saline (PBS) w/o calcium or magnesium (D8537, Sigma) before adding 100 μL of experimental media containing compounds and nutrients at indicated concentrations to each well. Proliferation was measured using the Incucyte platform (Sartorius). Briefly, images were captured every 2-4 hours and automatically counted using the Incucyte Cell-by-Cell software analysis package. Proliferation counts were normalized to the first cell count obtained immediately after beginning experimental conditions. Viability was assessed at 48 hour time points for AsPC1 and BxPC3 cells and 24 hour timepoints for Panc1 and MiaPaCa2 cells using AlamarBlue (Fisher Scientific, A50100) according to the manufacturer’s instructions. Briefly, 10 μL of AlamarBlue was added to each well containing 100 μL of experimental media. Plates were gently agitated to promote mixing, then incubated under standard conditions for a fixed time interval. Fluorescence was measured using 545 nm excitation and 590 nm emission using a BioTek Synergy H1 fluorescent plate reader. For the trypan blue viability assay, 50,000 cells/well were plated on 6-well plates overnight and switched to experimental media as described previously. At the indicated time point cells were washed twice with 1 mL of PBS w/o calcium or magnesium (D8537, Sigma-Aldrich) and removed from the plate using 200 μL of Trypsin-EDTA (0.25%) (2520056, Fisher Scientific) incubated for 5 minutes, and quenched with 200 μL of cell culture media. The suspended cell solution was mixed with equal parts of trypan blue 0.4% (15250061, Fisher Scientific) and counted using the Cellometer Auto T4 (Nexcelom Bioscience) for cell counting and viability measurement.

### Cell culture reagents

The following is a list of chemical compounds used in cell culture experiments: 1 mM N-acetyl-L-cysteine (A9615, Sigma-Aldrich), 100 μM Trolox (238813, Sigma-Aldrich), 100 μM deferoxamine (D9533, Sigma-Aldrich), 5 μM Ferrostatin-1 (SML0583, Sigma-Aldrich), 50 μM Z-VAD-FMK (14463, Cayman Chemical?), 10 μM Necrostatin-1S (20924, Cayman Chemical), 0.16 – 50 μM IKE (27088, Cayman Chemical), 0.16 – 10 μM RSL3 (SML2234, Sigma-Aldrich), 25-100 μM TEPP-46 (HY-18657, MedChem Express), 10 μM compound 3k (36815, Cayman Chemical), 50 μM Malic enzyme inhibitor (HY-124861, MedChem Express), 8 and 32 mM DM-Malate (374318, Sigma-Aldrich), 8 mM DM-Succinate (73605, Sigma-Aldrich), 8 mM DM-α-ketoglutarate (28394, Cayman Chemical), 5 μM CB-839 (22038, Cayman Chemical), 300 μM BSO (14484, Cayman Chemical), 1 mM GSH-EE (14953, Cayman Chemical), and 0.1-0.5%DMSO (D4540, Sigma-Aldrich).

### Lipid and general ROS quantification

The cells were seeded 25,000 cells/well in 24 well plates overnight. Cells were then washed with PBS twice and incubated in the experimental media for 24 hours. Cells were then stained for 30 minutes with either 10 μM chloromethyl-2′, 7′-dichlorodihydrofuorescein diacetate (CM-H2DCFDA) (C6827, Fisher Scientific) for general ROS quantification or 10 μM C11-BODIPY (D3861, Invitrogen) for lipid peroxidation according to the manufacturers protocol. Cells were also stained for 30 minutes simultaneously with 1 μM 2′-[4-ethoxyphenyl]-5-[4-methyl-1-piperazinyl]−2,5′-bi-1H-benzimidazoletrihydrochloride trihydrate (Hoechst 33342, Fisher Scientific) for nuclear visualization and cell localization. Cells were then washed two times with PBS and placed in live imaging solution (A14281DJ, Fisher Scientific) for image capture. Brightfield images and fluorescence were measured using a Leica DMi8 microscope, a PE4000 LED light source, 20x and 40x objective, a DFC9000GT camera, DAPI, GFP, or TexasRed filter set, and LAS X imaging software. 3 images for each fluorescent channel were captured for each well. For image processing and quantification, the images were imported into the Fiji version of ImageJ (http://fiji.sc). Cells in each image were selected, background selected and normalized to cell area. Fluorescence of cells for each well was determined by the average fluorescent signal for the 3 images for that well. Relative fluorescence is shown in the figures relative to control media conditions. Images shown are representative of the images captured for each condition.

### Metabolomic profiling and stable isotope labeling

To quantify metabolites, each cell line was seeded 200,000 cells/well in triplicate (n=3) in 6-well plates with media as described in the cell culture methodology section until achieving approximately 80% confluency (approximately 24 hours). For stable isotope labeling, media was refreshed on the plates and incubated for 2 hours. Plates were then washed with PBS then switched to labeling media containing either 1 mM ^13^C_5_-glutamine (CLM-1822-H-PK, Cambridge Isotope Laboratories) or 5 mM ^13^C_1,2_-glucose (CLM-504-PK, Cambridge Isotope Laboratories) and 10% dialyzed FBS (F0392, Sigma-Aldrich). Samples were collected at 0 minutes (unlabeled control) and 4 hours for ^13^C_1,2_-glucose and ^13^C_5_-glutamine labeling for the Panc1 cells. AsPC1 samples were collected identically with the exception that ^13^C-glutamine labeling was also tested at 24 hours. Metabolite extraction was performed as described previously.^90^ Briefly, each well is washed with 0.9% saline (16005–092, VWR), then 500 μL of HPLC grade methanol is added followed by 300 μL of HPLC-grade water containing 0.5 μM camphorsulfonic acid as an internal control. Cells are scraped from the plate and the solution is transferred to a 1.5 mL Eppendorf tube containing 500 μL of HPLC-grade chloroform. Samples are vortexed for 10 minutes, then centrifuged at 4 °C 16,000xg for 15 minutes. The polar layer is then removed and dried by lyophilization. In addition to intracellular metabolites, 100 μL media samples were also taken from each well at the same time points and extracted using the same procedure outline above. Protein extracted from the cells was dissolved in 0.2 M potassium hydroxide aqueous solution overnight and quantified using Pierce BCA Protein Assay Kit (PI23225, Fisher Scientific). Extracted metabolites are then resuspended in HPLC-grade water containing 5 μM 1,4-piperazinediethanesulfonic acid (PIPES; P6757, Sigma-Aldrich) as an internal standard.

LC-MS/MS analysis was performed with ion-pairing reverse phase chromatography using an Ascentis Express column (C18, 5 cm × 2.1 mm, 2.7 μm, Sigma-Aldrich) for separation and a Waters Xevo TQ-XS triple quadrupole mass spectrometer. Metabolite peak processing was performed in MAVEN.^91,92^ Total metabolite abundance (labeled and unlabeled) was scaled by the camphorsulphonic acid internal standard and protein content. The entire dataset for each independent experiment was then normalized by Probabilistic Quotient Normalization.^93^ For isotope labeled samples, the data was corrected for the natural ^13^C abundance using IsoCor and reported as the abundance of labeled metabolite relative to the control cell under control media conditions.^94^

### Pyruvate Kinase Activity Assay

Cells were plated 200,000 cells/well in 6-well plates and incubated overnight. Plates were washed twice with PBS w/o calcium or magnesium (Sigma, D8537) and the media was replaced with experimental media in triplicate for each condition as indicated. Panc1 cells were incubated for 12 hours, AspC1 were incubated for 24 hours as described above. Pyruvate kinase activity kit (MAK072, Sigma-Aldrich) was used according to the manufacturer’s protocol to evaluate pyruvate kinase activity. Briefly cells were lysed and scraped from the plate and centrifuged at 16,000xg for 5 minutes to clear debris. 5 μL of supernatant was used for each sample for the assay. The Pierce BCA Protein Assay Kit (PI23225, Fisher Scientific) was used to quantify protein extracted from the cells to normalize activity. After quantifying PK activity in nmole/min/mL, activity was normalized to the control cell line under normal media conditions to show relative PK activity.

### NADPH Quantification

Cells were plated in a white 96-well plate with clear bottoms at 8,000 cells/well for each cell line tested. Cells were allowed to seed overnight before switching the media to experimental conditions for a 24 hour incubation in Panc1 and a 48 hour incubation in AsPC1. The media was aspirated from the wells, the cells were washed twice with PBS, and 50 μL of PBS was added to each well. The manufacturer’s protocol was then followed for the assay (G9081, Promega). Briefly, 50 μL of a 1% DTAB and bicarbonate base buffer were added to each sample and 50 μL of sample were moved to a new set of empty wells. The plate is then heated at 60°C for 15 minutes. After the plate has been heated, the plate incubates at room temperature for 10 minutes. To the base-treated samples, 50 μL of a solution containing equal parts of 0.4N HCl and Trizma base were added to each well. Luciferase detection reagent was prepared and 100 μL were added to each well. The plate was briefly mixed on a plate shaker and covered in aluminum foil to keep light from entering the reaction. The plate was incubated for 30-60 minutes at room temperature before the luminescence was read on a BioTek Synergy HI plate reader.

### *In vivo* PDAC xenograft studies

To inoculate mice and produce xenograft tumors in mice, AsPC1 Control and PKM2KO Clone #1 cells were grown and prepared into a solution of 5x10^7^ cells/mL in PBS w/o calcium or magnesium (Sigma, D8537). 100 μL of each solution (5x10^6^ cells) was injected subcutaneously into the right flank of 8 week old male NOD scid gamma (NSG) mice. 12 mice received the control cells, and 12 mice received the PKM2KO cells. After 1 month the two groups of tumor bearing mice were each divided into 2 groups of 6 mice to receive intraperitoneal injections of either 200 μL of 50 mg/kg IKE in 65% DW5 (5% Dextrose in water), 5% TWEEN-80, and 30% PEG-400 or 200 μL of vehicle control. Treatment was delivered 6 days a week for 2 total weeks. Tumor volume and mouse body weight were measured 4 times a week. Tumor volume was estimated by caliper measurement across the long and short axes of the tumor using the equation length * width^2^ * 0.5 = volume (mm^3^). After two weeks, the mice were euthanized, and tumors were excised.

To evaluate the combination of IKE and TEPP-46 *in vivo*. Panc1 WT cells were grown and prepared into a solution of 5x10^7^ cells/mL in PBS w/o calcium or magnesium (Sigma, D8537). 100 μL of each solution (5x10^6^ cells) was injected subcutaneously into the right flank of 6 week old female NSG mice. After ∼3 weeks the tumor bearing mice were divided randomly into 4 groups (n=6 for each group except the IKE and TEPP46 combination which had n=4). Mice received daily intraperitoneal injections for 2 weeks of either 50 mg/kg IKE, 30 mg/kg TEPP-46, the combination of 50 mg/kg IKE with 30 mg/kg TEPP-46, or vehicle control. IKE treatment was delivered in 200 μL volumes and was formulated in 65% DW5 (5% Dextrose in water), 5% TWEEN-80, and 30% PEG300. TEPP-46 was delivered in 200 μL volumes and was formulated in 5% DMSO, 40% PEG300, 5% TWEEN-80, and 50% saline (0.9%). Tumor volume and mouse body weight were measured 4 times a week. Tumor volume was estimated by caliper measurement across the long and short axes of the tumor using the equation length * width^2^ * 0.5 = volume (mm^3^). After two weeks, the mice were euthanized, and tumors were excised.

### Statistical Analysis and Graph Generation

All data are expressed as means +/- standard deviation. Each experiment was conducted with three to six replicates and was reproduced in at least two independent experiments unless otherwise indicated. The figures shown are the results of one independent experiment representative of the results reproducible in each independent experiment. Statistical analysis and graphing were performed in Graph Pad v10 using Student’s T-Test, one-Way ANOVA, or two-Way ANOVA and corrected for multiple hypothesis testing using the Tukey test or Sidak test where appropriate. A *p*-value of < 0.05 was used as cutoff for determining significance. In figures * or # = *p*-val <0.05, ** = *p*-val < .01, *** = *p*-val < 0.001, **** = *p*-val <0.0001.

## Data Availability

All data supporting the findings of this study are available from the corresponding author on request.

## Acknowledgements

Support from NIH under award numbers R01CA270136 and R01AI139074 and the Aitch Foundation. Overview schematics were made with BioRender. We thank the MSU Mass spectrometry core facility and the additional present and former members of the Lunt lab: Dr. Deanna Broadwater, Amir Roshanzadeh, Dr. Shao Thing Teoh, and Dr. Martin Ogrodzinski for helpful suggestions.

## Author Contributions

E.E.: Conceptualization, investigation, methodology, formal analysis, and writing. T.J.: methodology, formal analysis, and writing – review and editing. H.M.: methodology, and writing – review and editing. G.T.: methodology and writing – review and editing. A.P.: methodology and writing – review and editing. L.Y.: methodology and writing – review and editing. S.Y.L.: conceptualization, writing—review and editing, supervision, project administration, and funding acquisition.

## Ethics

The authors declare no competing interests.

## Abbreviations

μL: Microliter
μm: Micrometer
μM: Micromolar
Arg: Arginine
BSO: Buthionine sulfoximine
Cas9: Caspase 9
CRISPR: Clustered regularly interspaced short palindromic repeats
Cys: Cystine
DFO: Deferoxamine
DMEM: Dulbecco’s modified eagle medium
DM-Mal: Dimethyl malate
DM-Suc: Dimethyl succinate
DM-αkg: Dimethyl α-ketoglutarate
FER: Ferrostatin-1
Fig.: Figure
Gln: Glutamine
Glu: Glutamate
Gly: Glycine
GOT1: Glutamate oxaloacetate transaminase 1
GOT2: Glutamate oxaloacetate transaminase 2
GPX4: Glutathione peroxidase 4
GSH: Reduced glutathione
GSH-EE: Reduced glutathione ethyl ester
GSSG: Oxidized glutathione
H2DCFDA: 2’,7’-dichlorodihydrofluorescein diacetate
His: Histidine
HPLC: High performance liquid chromatography
IKE: Imidazole ketone erastin
Ile: Isoleucine
KO: Knockout
LC-MS/MS: Liquid chromatography tandem mass spectrometry
Leu: Leucine
Lys: Lysine
M: mass
ME1: Malic enzyme 1
ME1i: Malic enzyme1 inhibitor
MEF: Mouse embryonic fibroblasts
Met: Methionine
ml: Milliliter
mM: Millimolar
N: Sample size
NAC: N-acetylcysteine
NEC: Necrostatin-1S
nm: nanometer
PDAC: Pancreatic ductal adenocarcinoma
NSG: NOD scid gamma
Phe: Phenylalanine
PK: Pyruvate kinase
PKM1: Pyruvate kinase muscle isoform 1
PKM2: Pyruvate kinase muscle isoform 2
PPP: Pentose phosphate pathway
p-val: p value
ROS: reactive oxygen species
RSL3: Ras selective lethal 3
Ser: Serine
TCA: Tricarboxylic acid
Thr: Threonine
TRO: Trolox
Trp: Tryptophan
Tyr: Tyrosine
U: Uniform
Val: Valine
Vec: Vector
WT: Wild-type
xCT: SLC7A11, component of system X_c_^−^ cystine/glutamate antiporter
ZVAD: Z-VAD-FMK

## Supplemental Figures

**Supplementary Figure S1.**
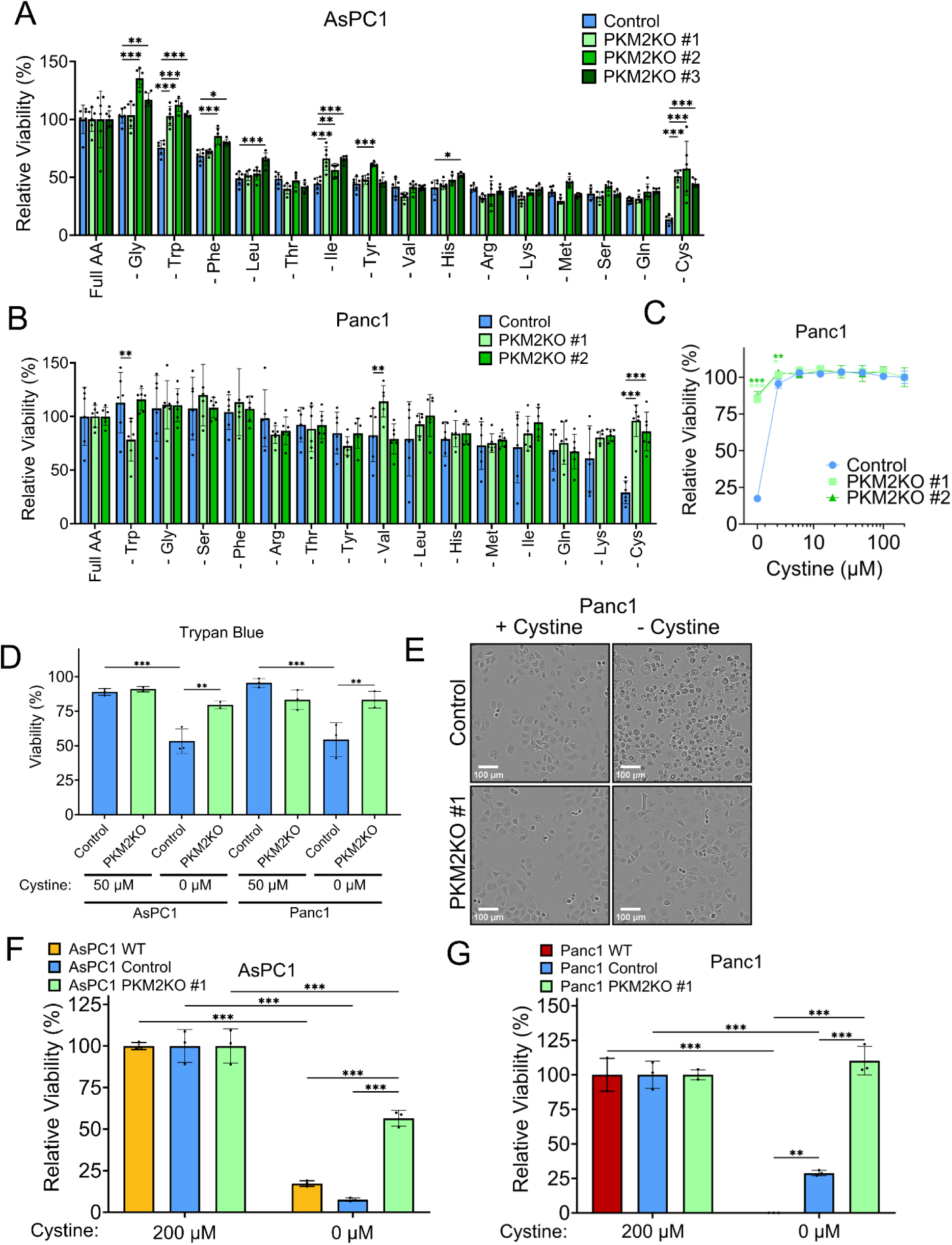
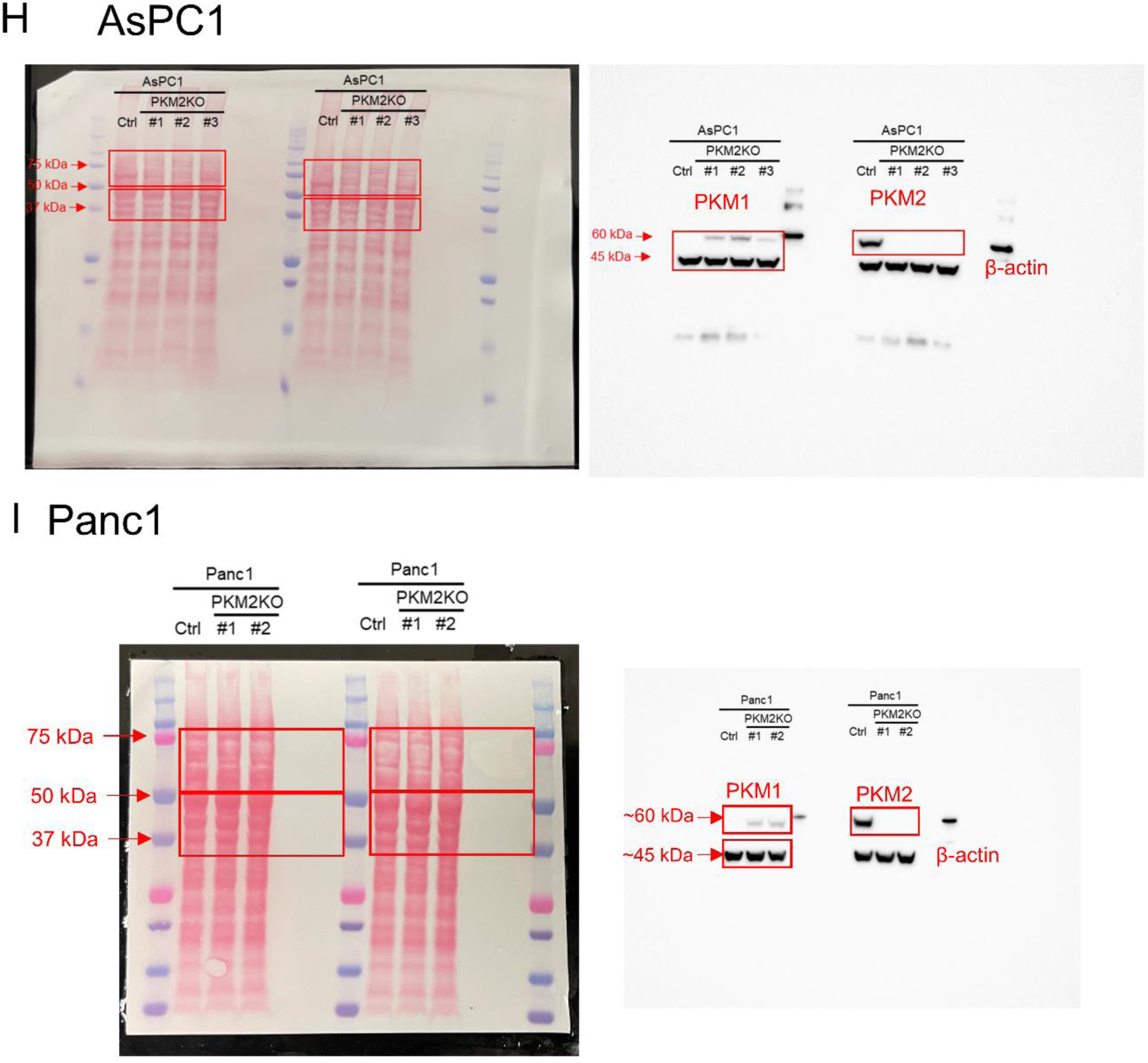
Additional data supporting Figure 1. **A-B.** Relative viabilities of AsPC1 (**A**) and Panc1 (**B**) control and respective PKM2KO clones in DMEM without each individual amino acid as shown. Significance was assessed by two-way ANOVA. \*\**p*<0.01, \*\*\**p*<0.001. Multiple hypothesis correction by Tukey test. **C.** Relative viabilities of Panc1 control and PKM2KO clones under a range of cystine concentrations from 200-0 μM. Significance was assessed by two-way ANOVA. \**p*<0.05, \*\**p*<0.01, \*\*\**p*<0.001. Multiple hypothesis correction by Dunnet test. **D.** Relative viabilities of AsPC1 and Panc1 control and PKM2KO cells under 50 μM and 0 μM cystine determined using trypan blue. Significance was assessed by two-way ANOVA. \**p*<0.05, \*\**p*<0.01, \*\*\**p*<0.001. Multiple hypothesis correction by Tukey test. **E.** Brightfield microscopy images of Panc1 Control and PKM2KO cells under either 200 or 0 μM cystine conditions. Scale bar = 100 μm. **F, G.** Relative viabilities of AsPC1 (**F**) and Panc1 (**G**) WT, control, and PKM2KO cells under 200 and 0 μM cystine. Significance was assessed by two-way ANOVA. \**p*<0.05, \*\**p*<0.01, \*\*\**p*<0.001. Multiple hypothesis correction by Tukey test. **H-I.** Full images of Ponceau staining and western blot from cropped image in Figure 1B. Red boxes indicated cut regions of membrane for primary antibody incubation and cropped portions of immunoblot.

**Supplementary Figure S2.**
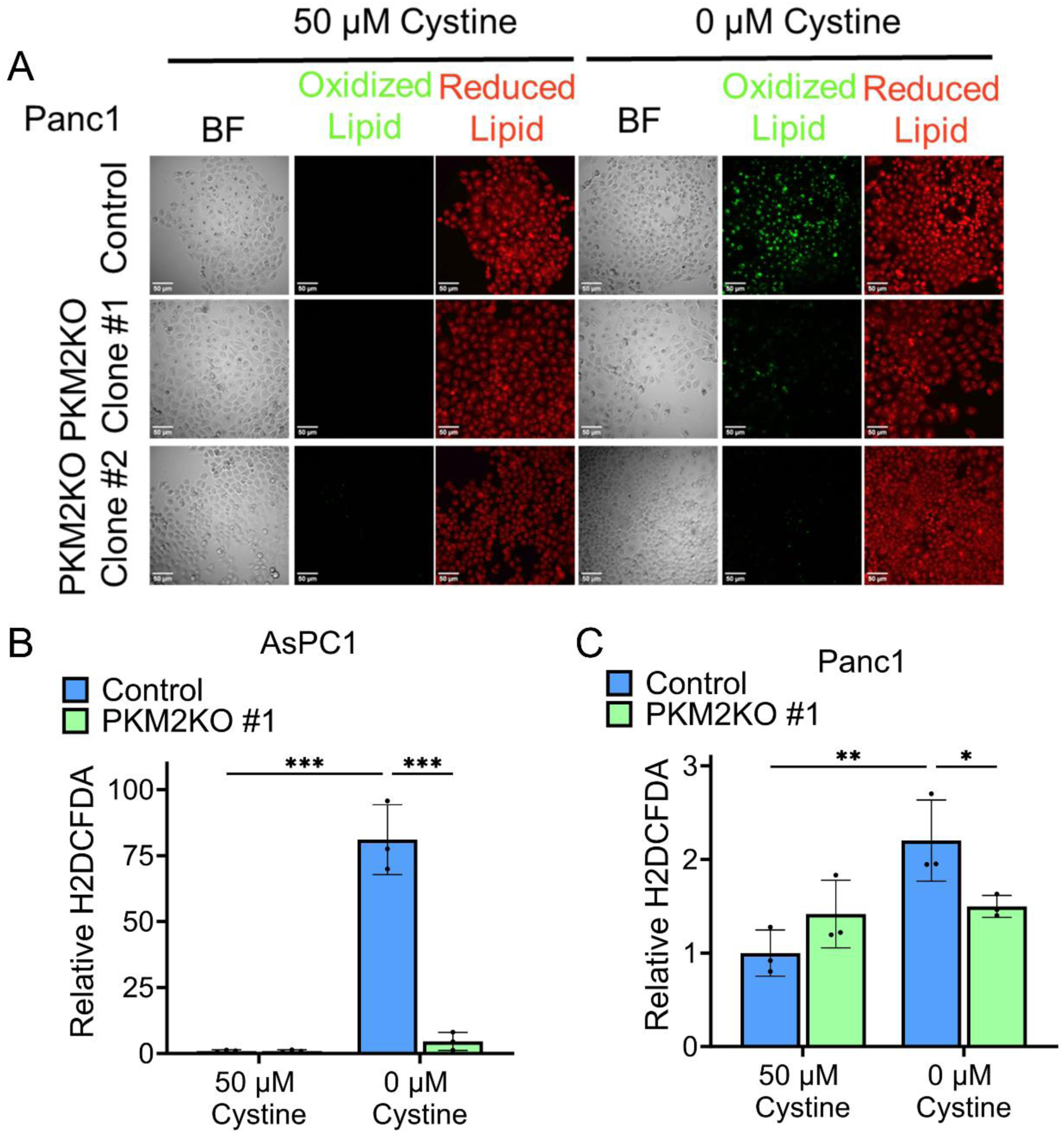
Additional data supporting figure 2. **A.** Representative brightfield and fluorescent images of Panc1 control and PKM2KO cell lipid peroxidation quantified in Fig. 2E. Scale bar = 50 μm. **B, C.** Relative general ROS production measured by H2DCFDA in AsPC1 (**B**) and Panc1 (**C**) control and PKM2KO cells under 50 and 0 μM cystine. Significance was assessed by two-way ANOVA. \**p*<0.05, \*\**p*<0.01, \*\*\**p*<0.001. Multiple hypothesis correction by Tukey test.

**Supplementary Figure S3.**
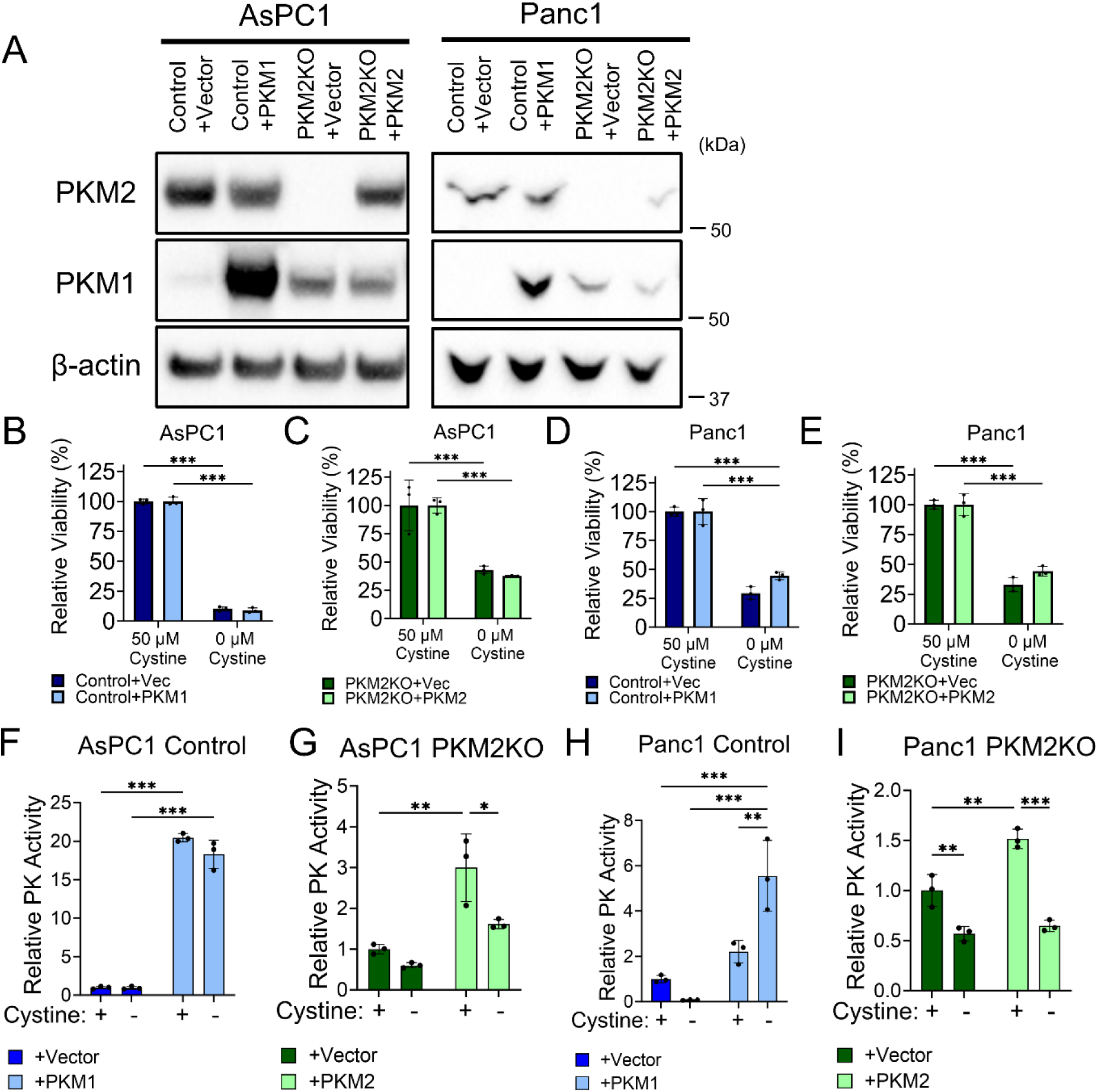
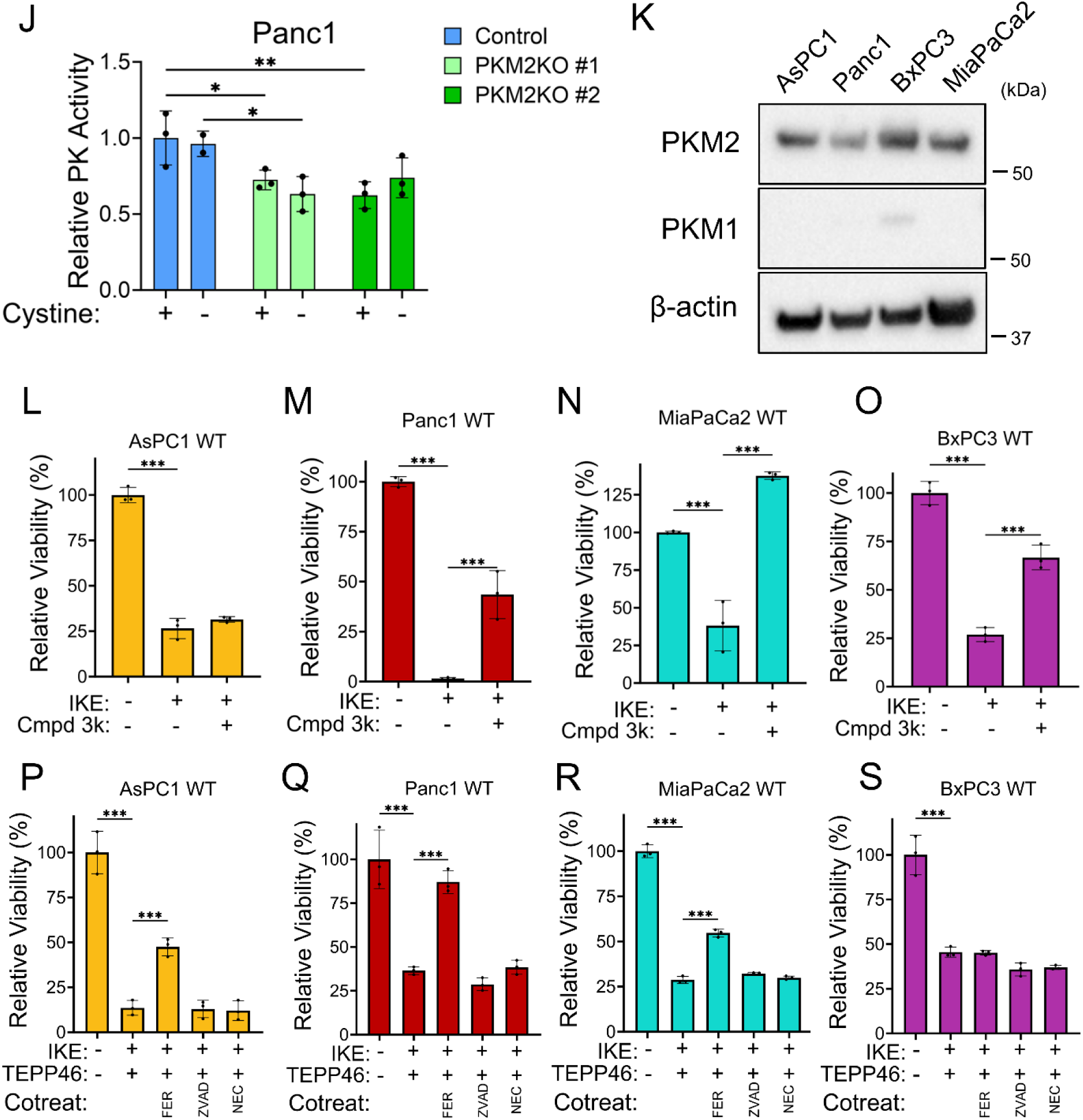
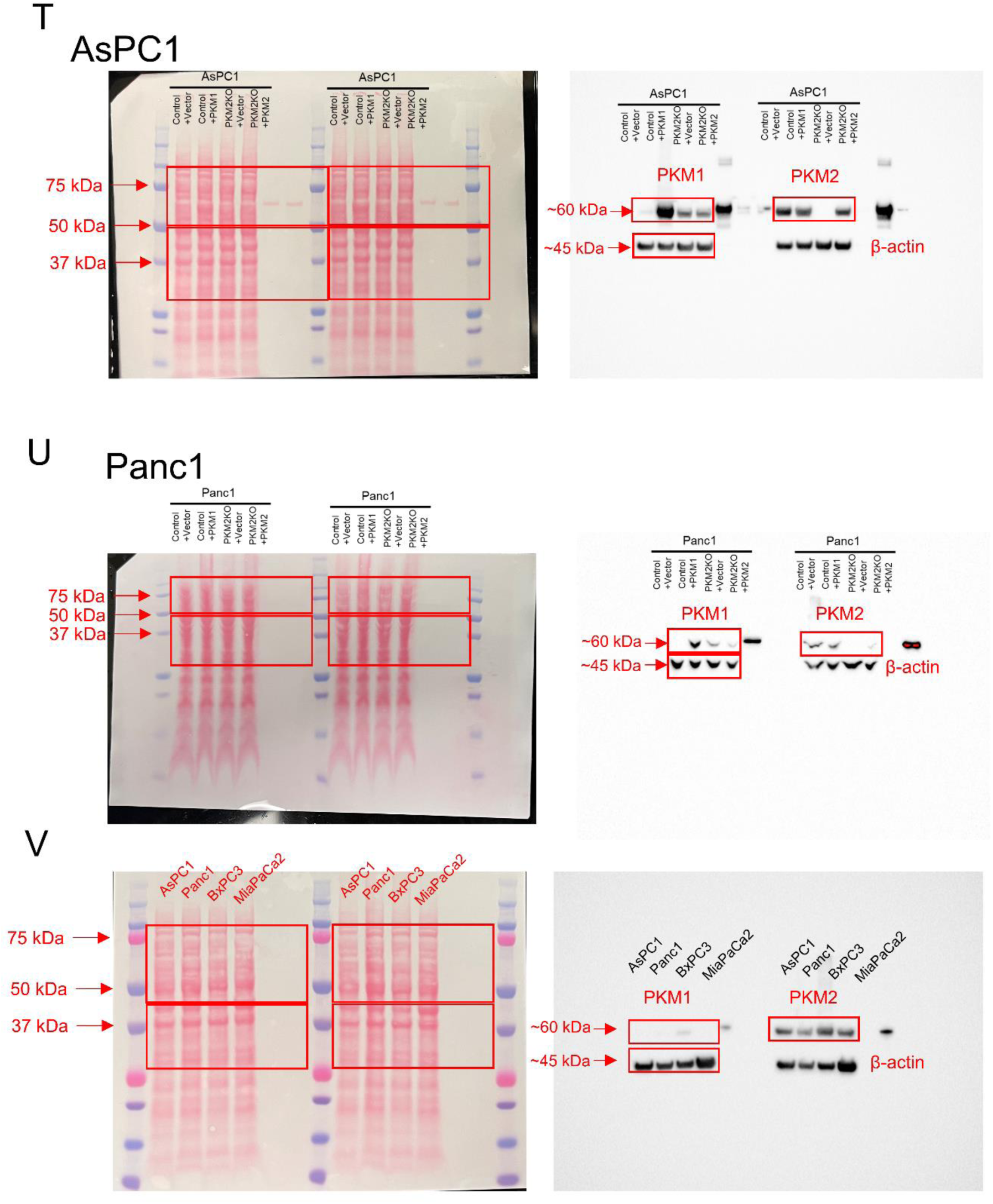
Additional data in support of figure 3. **A.** Western blot of PKM2 and PKM1 expression in AsPC1 and PKM2 control + vector, control + PKM1, PKM2KO + vector, and PKM2KO + PKM2. **B.** Relative viabilities of AsPC1 control + vector and control + PKM1 cells under 50 μM and 0 μM cystine. **C.** Relative viabilities of AsPC1 PKM2KO + vector and PKM2KO + PKM2 cells under 50 μM and 0 μM cystine. **D.** Relative viabilities of Panc1 control + vector and control + PKM1 cells under 50 μM and 0 μM cystine. **E.** Relative viabilities of Panc1 PKM2KO + vector and PKM2KO + PKM2 cells under 50 μM and 0 μM cystine. For **B**-**E**, significance was assessed by two-way ANOVA. \**p*<0.05, \*\**p*<0.01, \*\*\**p*<0.001. Multiple hypothesis correction by Tukey test. **F.** Relative pyruvate kinase activity of AsPC1 control + vector and control + PKM1 cells under 50 μM (+) and 0 μM (-) cystine. **I.** Relative pyruvate kinase activity of AsPC1 PKM2KO + vector and PKM2KO + PKM2 cells under 50 μM (+) and 0 μM (-) cystine. **J.** Relative pyruvate kinase activity of Panc1 control + vector and control + PKM1 cells under 50 μM (+) and 0 μM (-) cystine. **K.** Relative pyruvate kinase activity of Panc1 PKM2KO + vector and PKM2KO + PKM2 cells under 50 μM (+) and 0 μM (-) cystine. For **F**-**I**, Significance was assessed by two-way ANOVA. \**p*<0.05, \*\**p*<0.01, \*\*\**p*<0.001. Multiple hypothesis correction by Tukey test. **J.** Relative pyruvate kinase activity in Panc1 control and PKM2KO cells under 50 and 0 μM cystine. Significance was assessed by two-way ANOVA. \**p*<0.05, \*\**p*<0.01. Multiple hypothesis correction by Tukey test. **K.** Western blot of PKM1 and PKM2 expression in AsPC1, Panc1, BxPC3, and MiaPaCa2 WT cells. **L.** Relative viability of AsPC1 WT cells treated with (+) or without (-) 2.5 μM compound 3k and 5 μM IKE. **M.** Relative viability of Panc1 WT cells treated with (+) or without (-) 10 μM compound 3k and 10 μM IKE. **N.** Relative viability of MiaPaCa2 WT cells treated with (+) or without (-) 10 μM compound 3k and 5 μM IKE. **O.** Relative viability of BxPC3 WT cells treated with (+) or without (-) 2.5 μM compound 3k and 5 μM IKE.**P-S.** Relative viabilities of AsPC1 WT cells (**P**) and BxPC3 WT cells (**S**) treated with (+) or without (-) 10 μM IKE and 50 μM TEPP-46, and Panc1 WT cells (**Q**) and MiaPaCa2 WT cells (**R**) treated with (+) or without (-) 10 μM IKE and 50 μM TEPP-46. In **P-S** cells each were co-treated with 5 μM ferrostatin-1 (FER), 50 μM Z-VAD-FMK (ZVAD), 10 μM necrostatin-1S (NEC). For **L**-**S**, significance was assessed by two-way ANOVA. \**p*<0.05, \*\**p*<0.01, \*\*\**p*<0.001. Multiple hypothesis correction by Tukey test. **T-U.** Full images of Ponceau staining and western blot from cropped image in Figure S3A. **V.** Full images of Ponceau staining and western blot from cropped image in Figure S3O. For **T-V,** Red boxes indicate cut regions of membrane for primary antibody incubation and cropped portions of immunoblot.

**Supplementary Figure S4.**
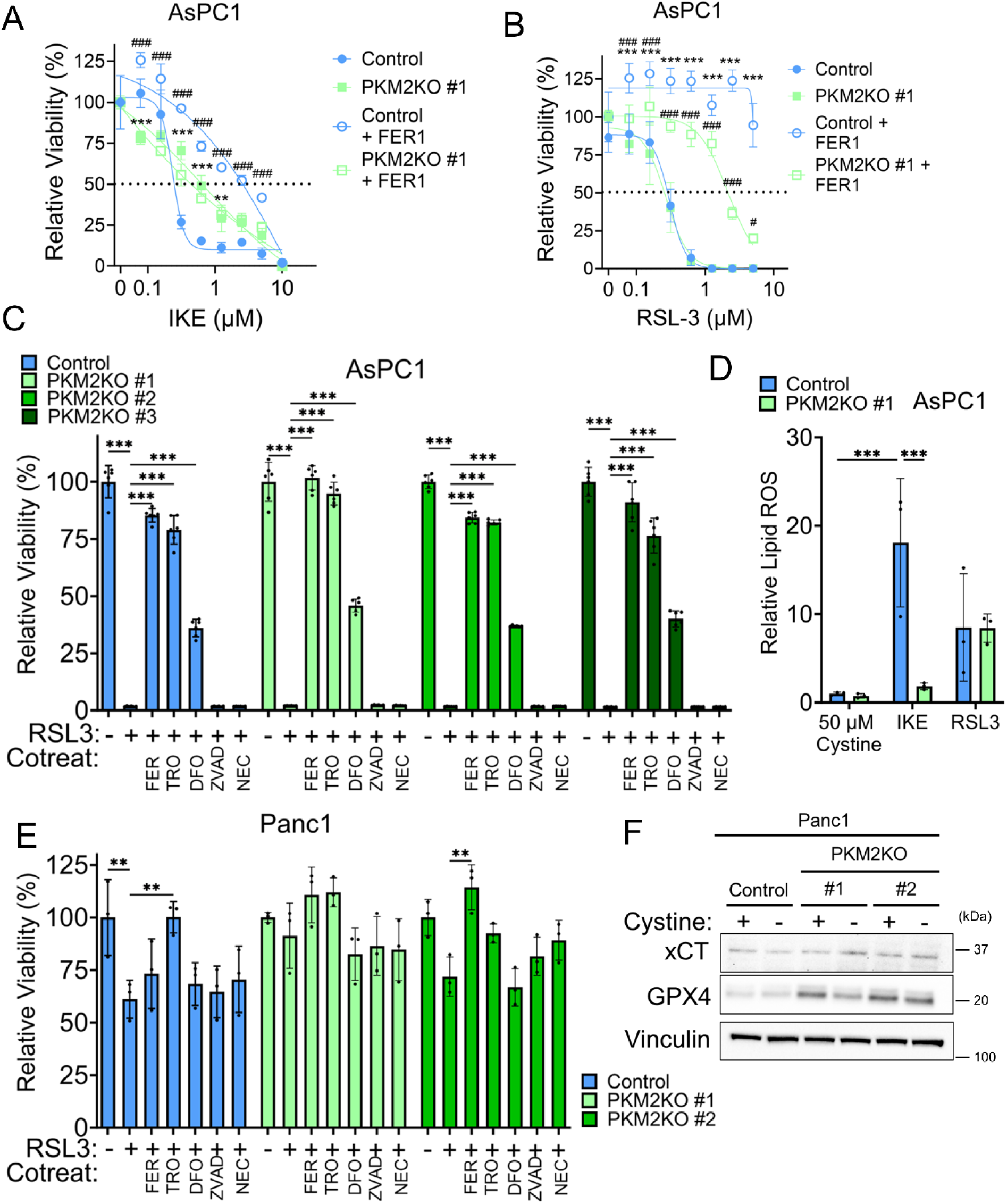
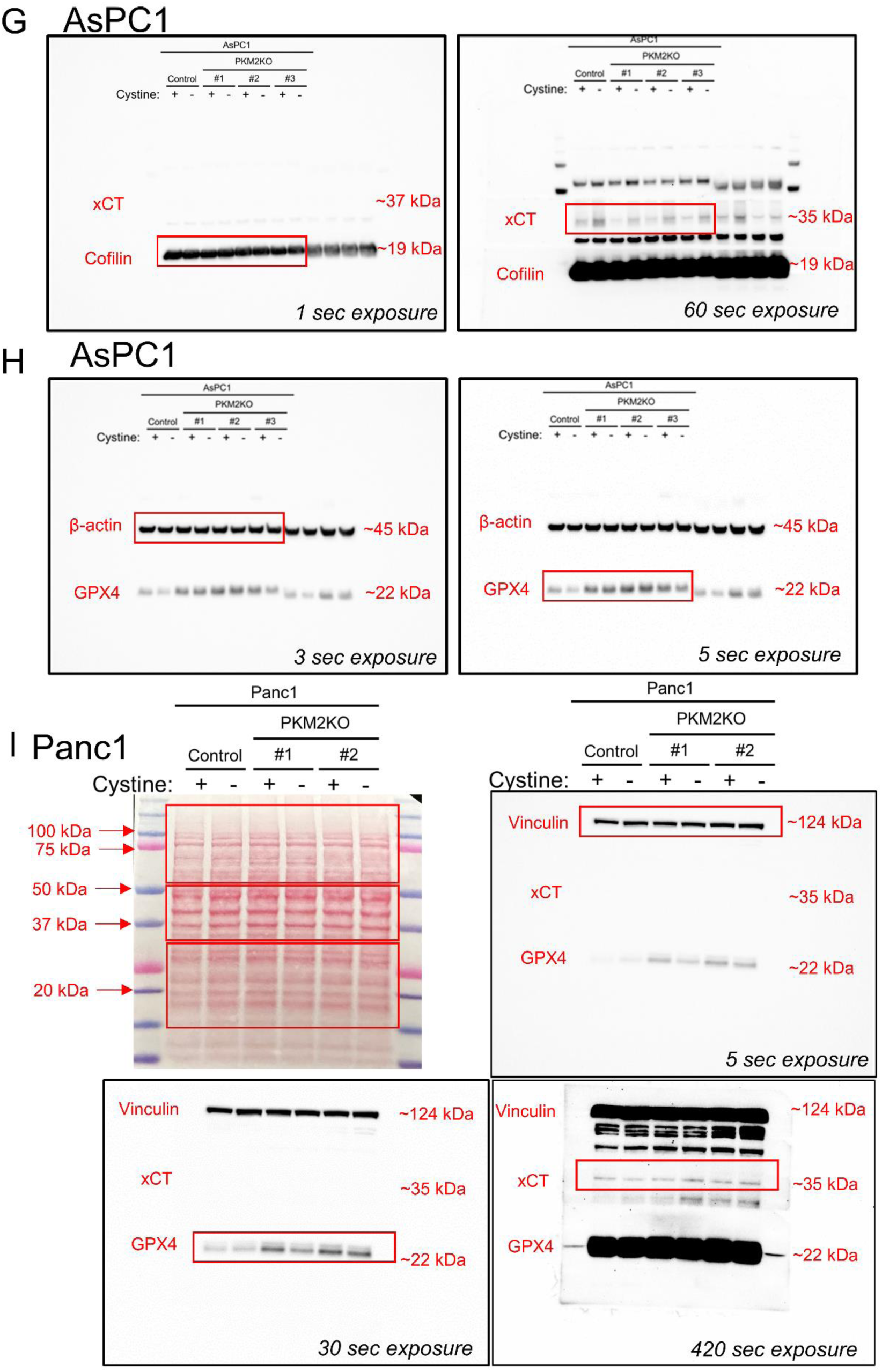
Additional data in support of figure 4. **A.** Concentration dependent viability responses of AsPC1 control and PKM2KO clone #1 to a range of IKE concentrations from 10-0 μM and co-treated with 5 μM ferrostatin-1 (FER1). Significance was assessed by two-way ANOVA. Between control and PKM2KO without FER1, \*\**p*<0.01, \*\*\**p*<0.001. Between control and control with FER1, ###*p*<0.001. Multiple hypothesis correction by Tukey test. **B.** Concentration dependent viability responses of AsPC1 control and PKM2KO clone #1 to a range of RSL3 concentrations from 5-0 μM and co-treated with 5 μM Ferrostatin-1 (FER). Significance was assessed by two-way ANOVA. Between control and control with FER1, \*\*\**p*<0.001. Between PKM2KO and PKM2KO with FER1, #*p*<0.05, ###*p*<0.001. Multiple hypothesis correction by Tukey test. **C, E.** Relative viabilities of AsPC1 (**C**) and Panc1 (**E**) control and PKM2KO cells under 50 μM cystine and 5 μM RSL3 co-treated with 5 μM ferrostatin-1 (FER), 100 μM trolox (TRO), 100 μM deferoxamine (DFO), 50 μM Z-VAD-FMK (ZVAD), 10 μM necrostatin-1S (NEC). Significance was assessed by two-way ANOVA. \**p*<0.05, \*\**p*<0.01, \*\*\**p*<0.001. Multiple hypothesis correction by Tukey test. **D.** Relative lipid peroxidation visualized by C11-BODIPY of AsPC1 control and PKM2KO #1 under 50 μM cystine, 5 μM IKE, and 5 μM RSL3 treatment. Significance was assessed by two-way ANOVA. \**p*<0.05, \*\**p*<0.01, \*\*\**p*<0.001. Multiple hypothesis correction by Tukey test. **F.** Western blot of xCT and GPX4 expression in Panc1 control and PKM2KO cells under 50 μM (+) and 0 μM (-) cystine. **G-H.** Full images of western blot from cropped image in Figure 4B **I.** Full images of Ponceau staining and western blot from cropped image in Figure S4F. Red boxes indicate cut regions of membrane for primary antibody incubation and cropped portions of immunoblot.

**Supplementary Figure S5.**
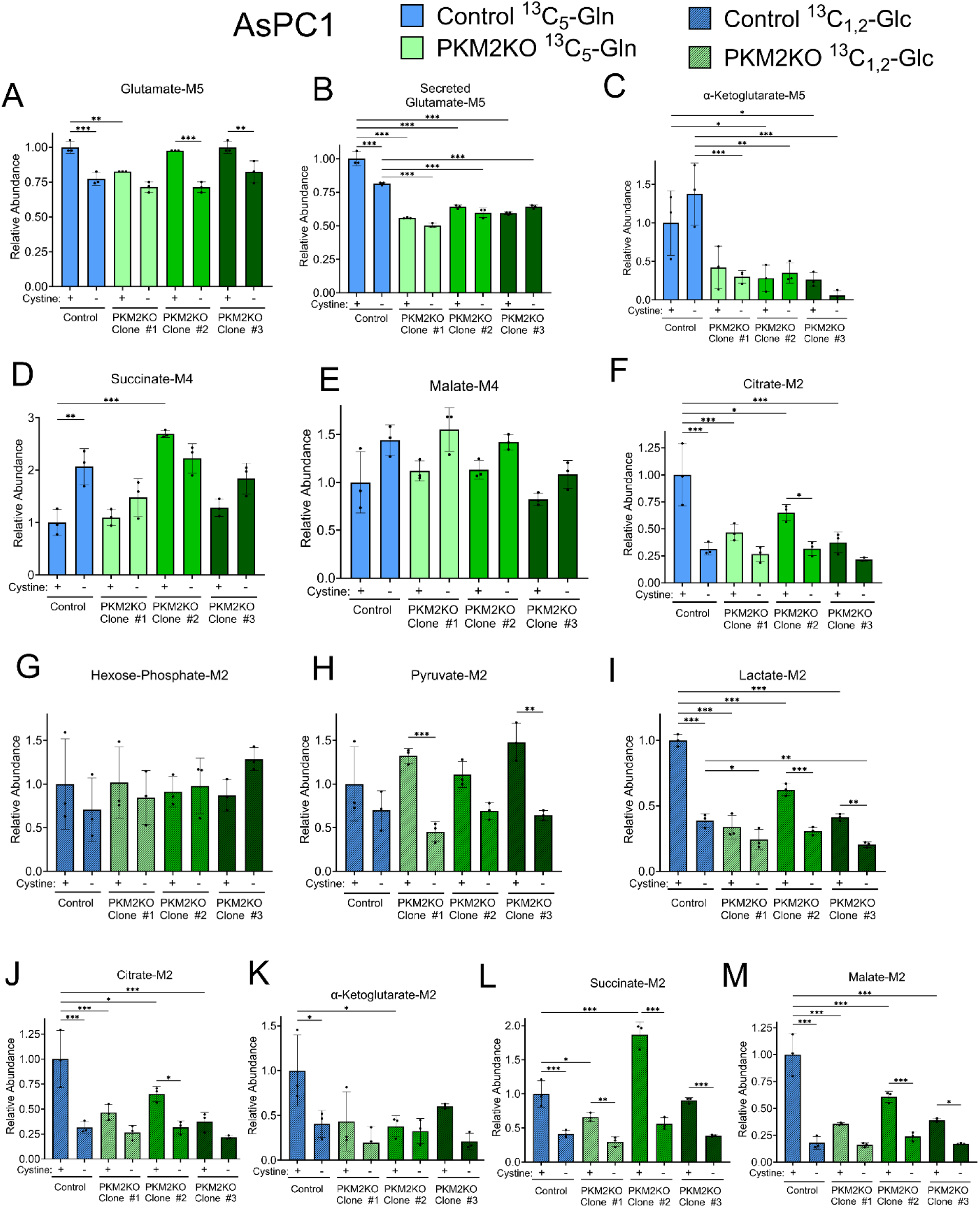
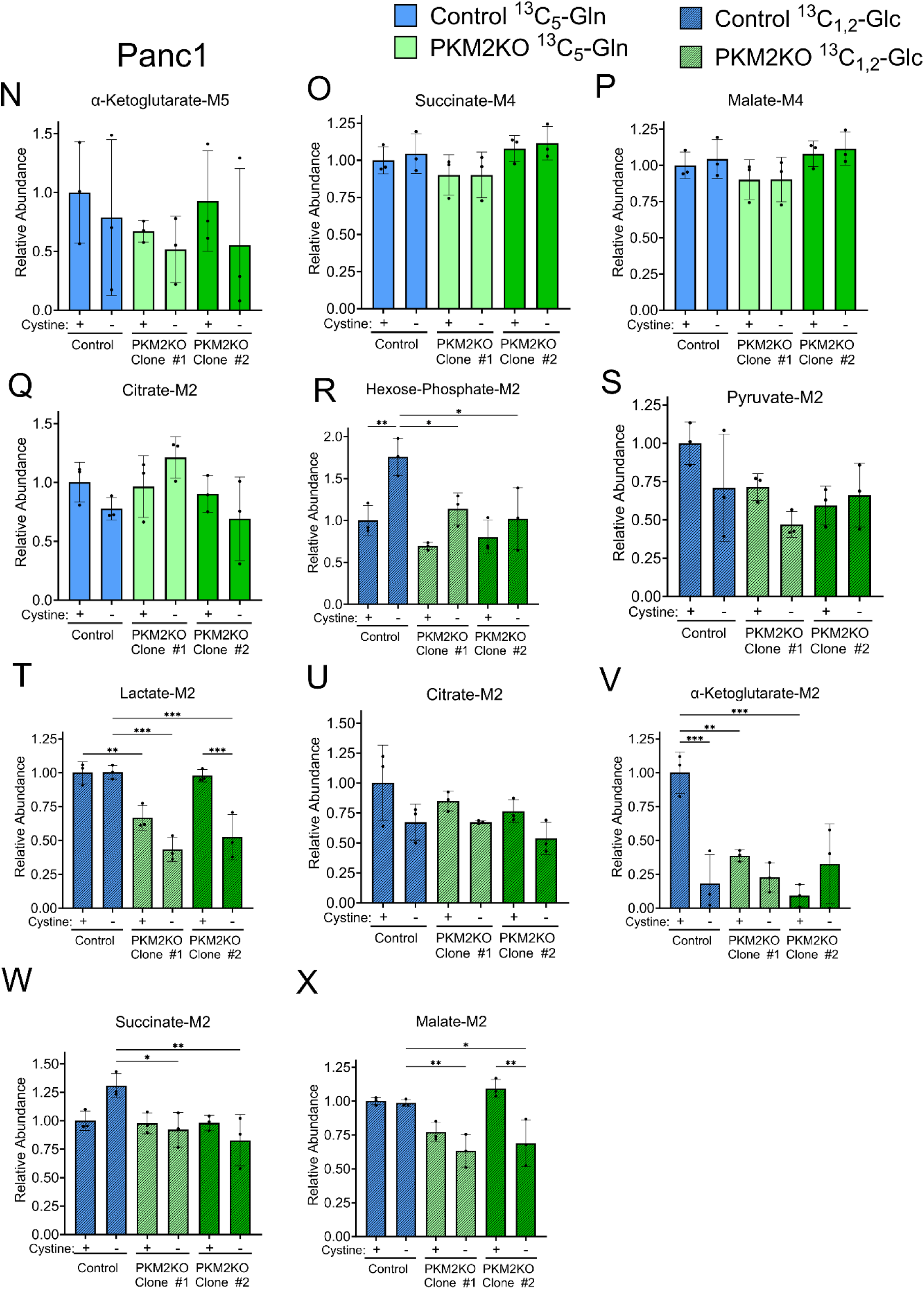
Additional data in support of Figure 5. **A.-H.** Relative abundance of indicated metabolites from stable isotope tracing of U-^13^C_5_-glutamine under 50 and 0 μM cystine for 24 hours in AsPC1 control and all PKM2KO clones. **G.-M.** Relative abundance of indicated metabolites from stable isotope tracing of ^13^C_1,2_-glucose under 50 μM (+) and 0 μM (-) cystine for 4 hours in AsPC1 control and all PKM2KO clones. **N.-Q.** Relative abundance of indicated metabolites from stable isotope tracing of U-^13^C_5_-glutamine under 50 μM (+) and 0 μM (-) cystine for 24 hours in AsPC1 control and all PKM2KO clones. **R.-X.** Relative abundance of indicated metabolites from stable isotope tracing of ^13^C_1,2_-glucose under 50 and 0 μM cystine for 4 hours in AsPC1 control and all PKM2KO clones. For **A**-**X**, significance was assessed by two-way ANOVA. \**p*<0.05, \*\**p*<0.01, \*\*\**p*<0.001. Multiple hypothesis correction by Tukey test.

**Supplementary Figure S6.**
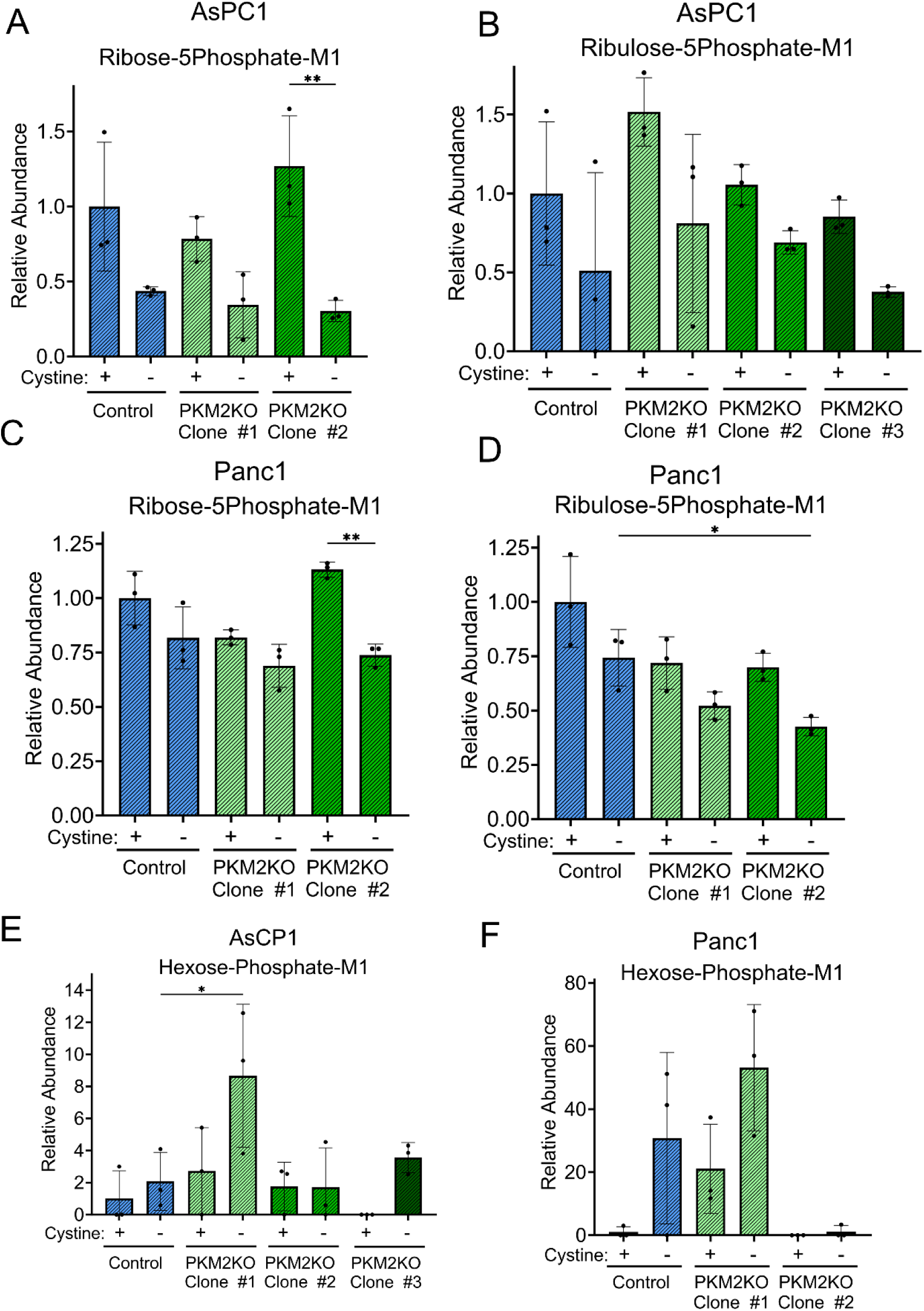
Glucose flux through the pentose phosphate pathway is unchanged in PKM2KO cells. **A**-**B**. Stable isotope tracing of ^13^C_1,2_-glucose under 50 μM (+) and 0 μM (-) cystine for 4 hours in AsPC1 control and all PKM2KO clones to produce M+1 labeled ribose-5phosphate (**A**) and ribulose-5phosphate (**B**). **C**-**D**. Stable isotope tracing of ^13^C_1,2_-glucose under 50 μM (+) and 0 μM (-) cystine for 4 hours in Panc1 control and all PKM2KO clones to produce M+1 labeled ribose-5phosphate (**C**) and ribulose-5phosphate (**D**). **E**-**F**. Stable isotope tracing of ^13^C_1,2_-glucose under 50 μM (+) and 0 μM (-) cystine for 4 hours in AsPC1 (**E**) and Panc1 (**F**) control and all PKM2KO clones to produce M+1 labeled hexose-phosphate. Significance was assessed by two-way ANOVA. \**p*<0.05, \*\**p*<0.01, \*\*\**p*<0.001. Multiple hypothesis correction by Tukey test.

**Supplementary Figure S7.**
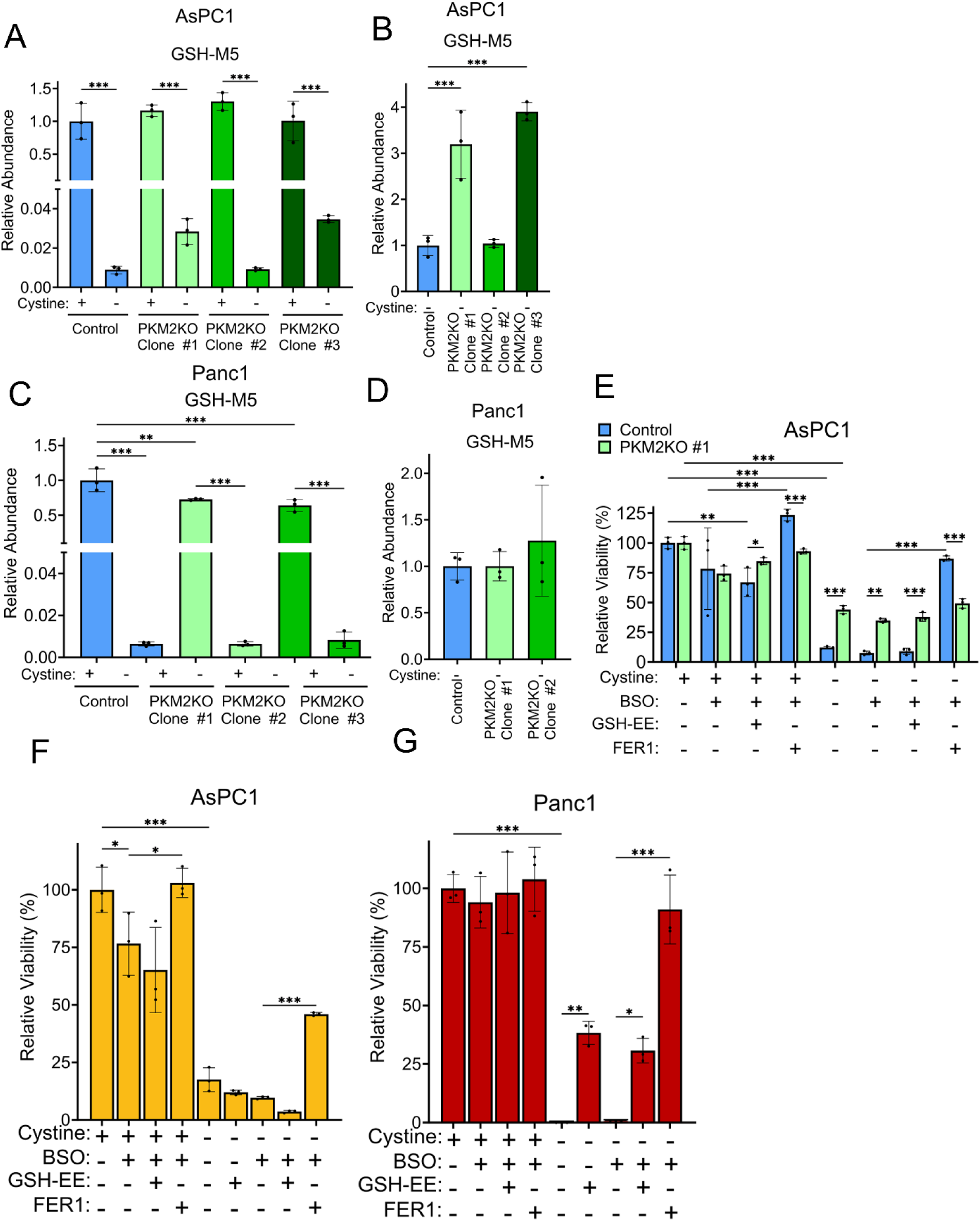
Glutathione synthesis is significantly increased in AsPC1 PKM2KO cells but is not essential for increasing viability under 0 μM cystine conditions. **A-B.** Stable isotope tracing of ^13^C_5_-glutamine under 50 μM (+) and 0 μM (-) cystine for 24 hours in AsPC1 control and all PKM2KO clones to produce M+5 labeled glutathione. **C.-D.** Stable isotope tracing of ^13^C_5_-glutamine under 50 μM (+) and 0 μM (-) cystine for 24 hours in Panc1 control and all PKM2KO clones to produce M+5 labeled glutathione. **E**-**G**. Relative viabilities of AsPC1 control and PKM2KO cells, AsPC1 WT cells (**F**), and Panc1 WT cells (**G**) under 50 μM (+) or 0 μM (-) cystine co-treated with 300 μM buthionine-sulfoximine (BSO), 1 mM glutathione-ethyl ester (GSH-EE), and/or 5 μM ferrostatin-1 (FER). Significance was assessed by two-way ANOVA.\**p*<0.05, \*\**p*<0.01, \*\*\**p*<0.001. Multiple hypothesis correction by Tukey test.

**Supplementary Figure S8.**
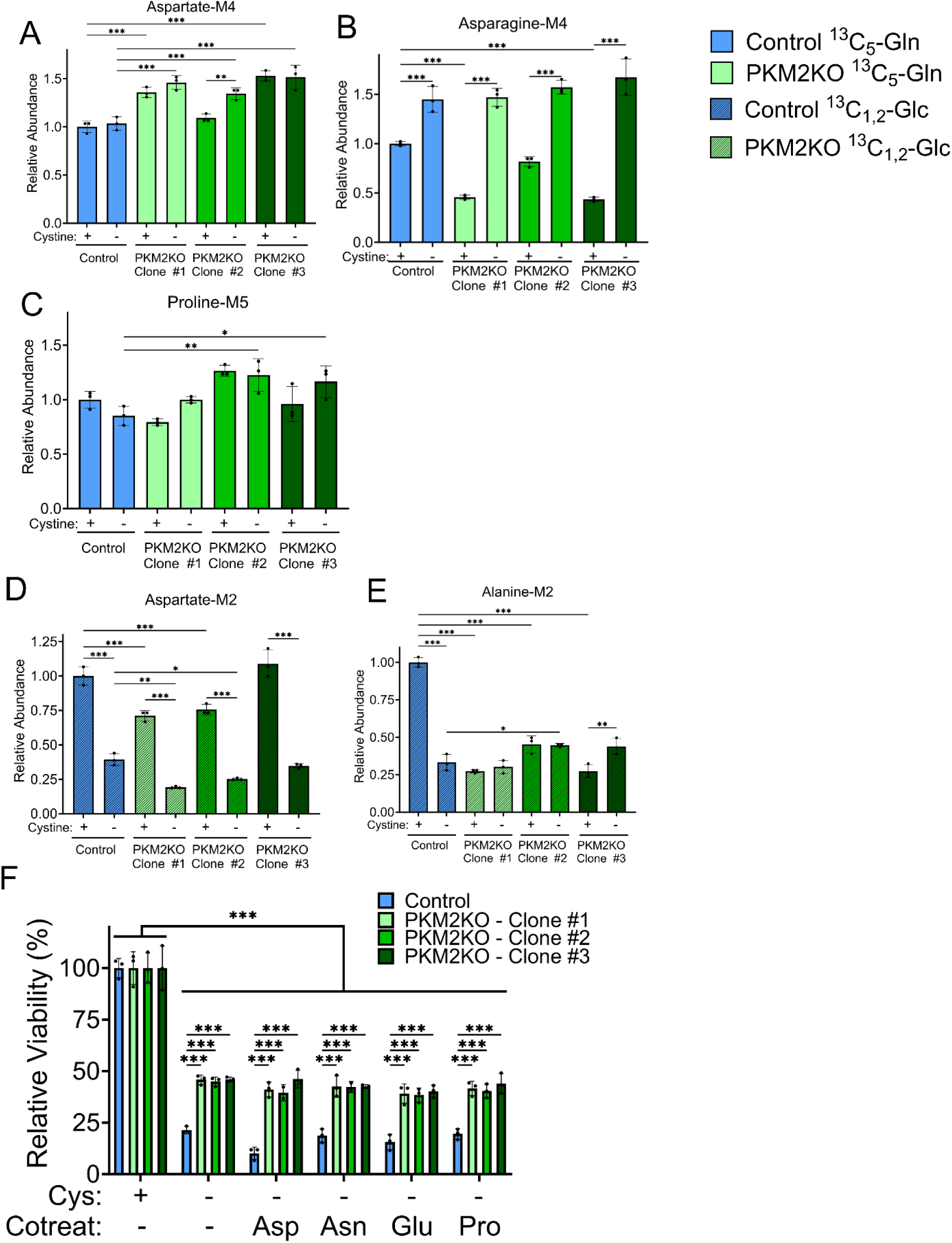
Amino acid synthesis from glutamine, but not glucose, is increased in PKM2KO cells under cystine starvation. **A-C**. Stable isotope tracing of ^13^C_5_-glutamine under 50 μM (+) and 0 μM (-) cystine for 24 hours in AsPC1 control and all PKM2KO clones to produce M+4 labeled aspartate (**A**) and asparagine (**B**), and M+5 proline (**C**). **D**-**E**. Stable isotope tracing of ^13^C_1,2_-glucose under 50 μM (+) and 0 μM (-) cystine for 4 hours in AsPC1 control and all PKM2KO clones to produce M+2 labeled aspartate (**D**) and alanine (**E**). **F**. Relative viabilities of AsPC1 control and PKM2KO cells under 50 μM (+) and 0 μM (-) cystine supplemented with either 6 μM aspartate, 90 μM asparagine, 90 μM glutamate, or 240 μM proline. Significance was assessed by two-way ANOVA. \**p*<0.05, \*\**p*<0.01, \*\*\**p*<0.001. Multiple hypothesis correction by Tukey test.

**Supplementary Figure S9.**
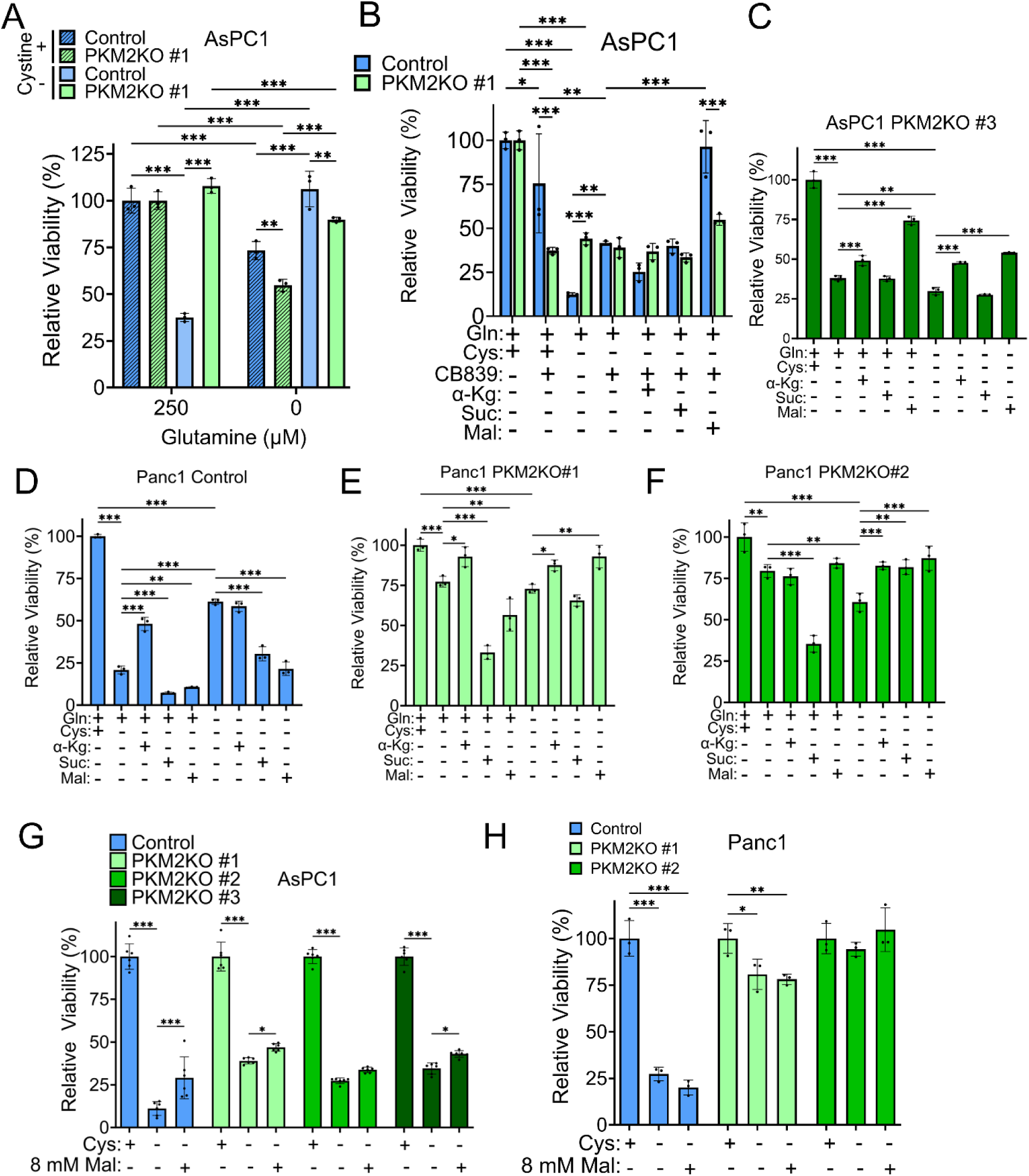
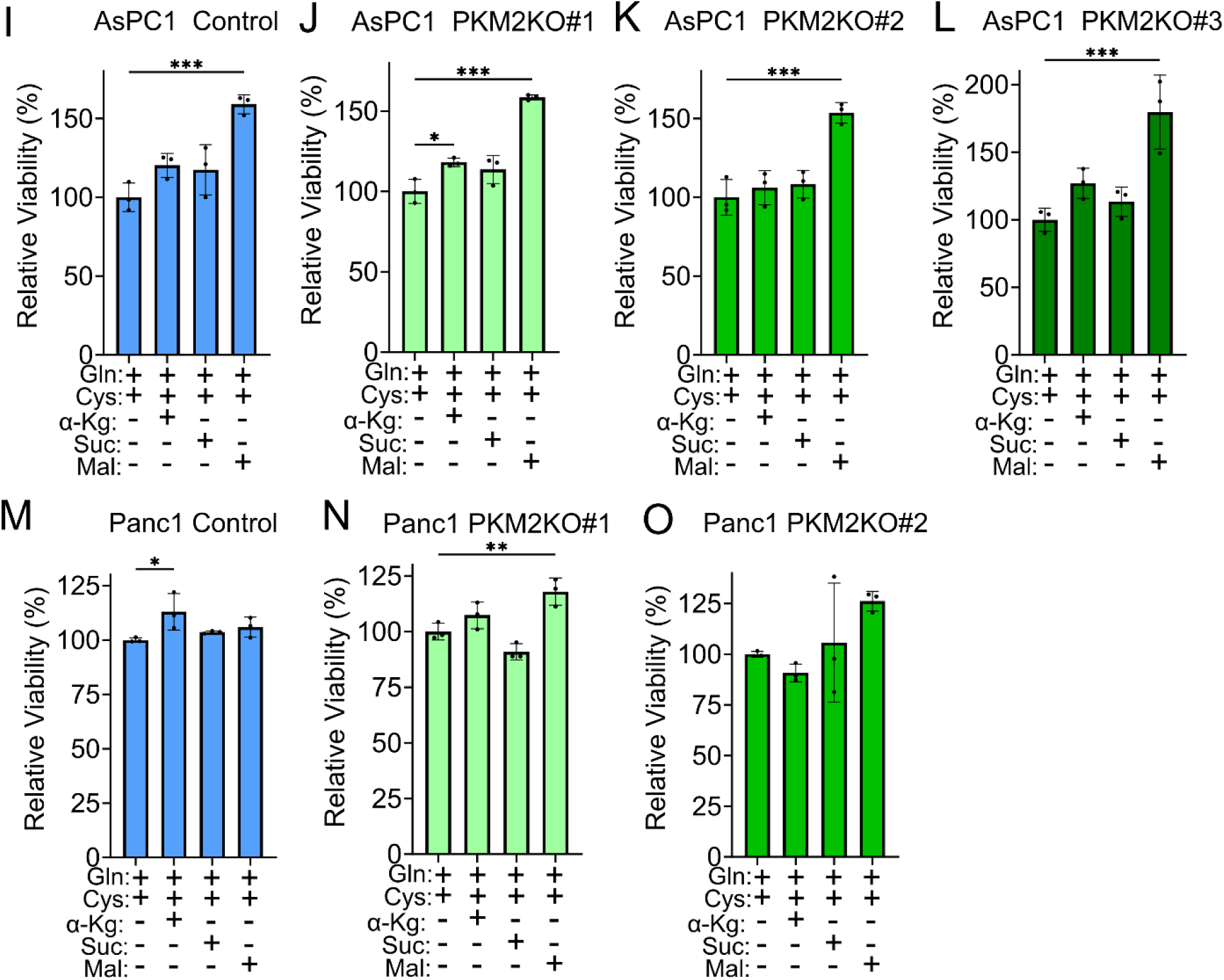
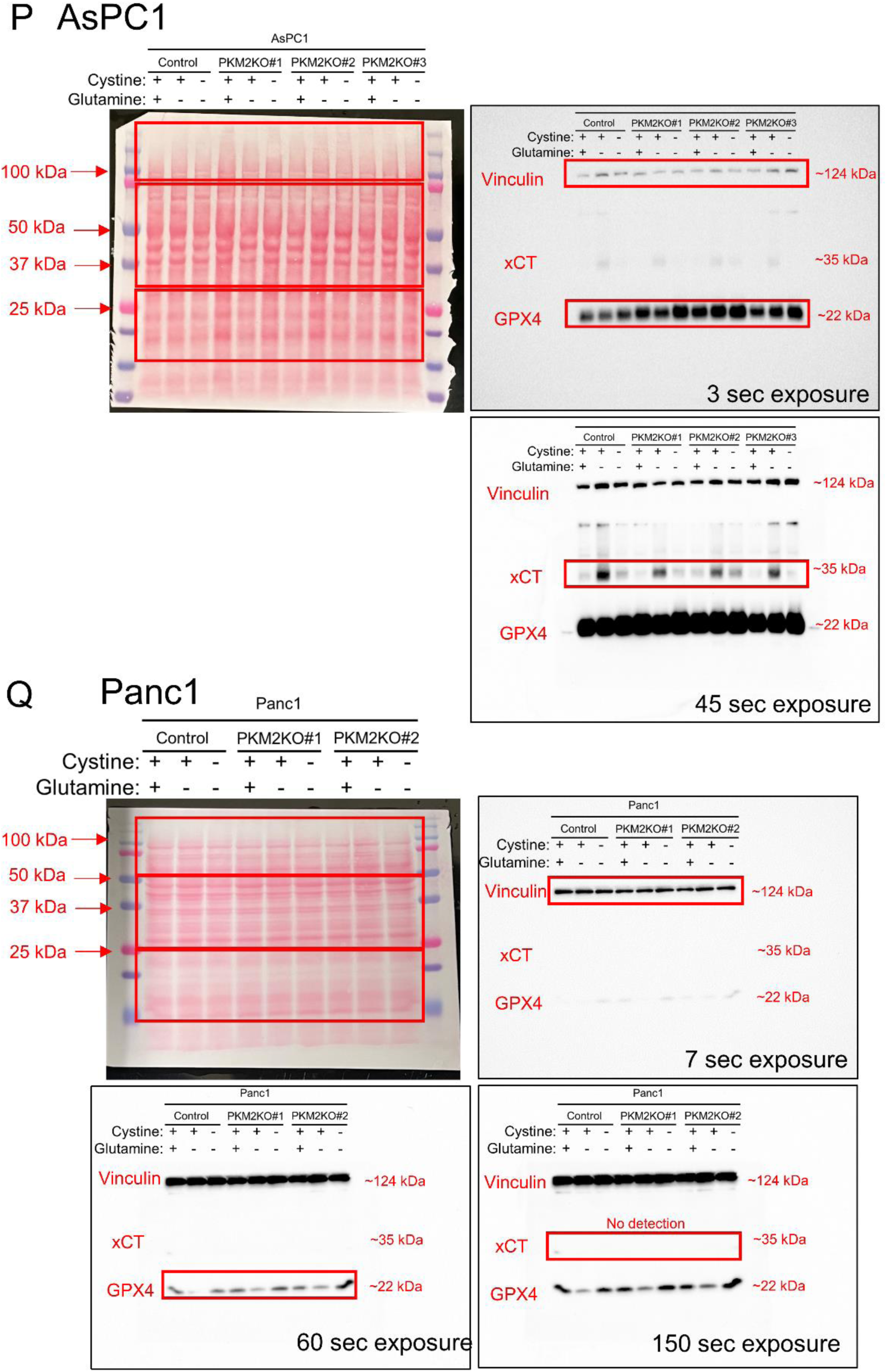
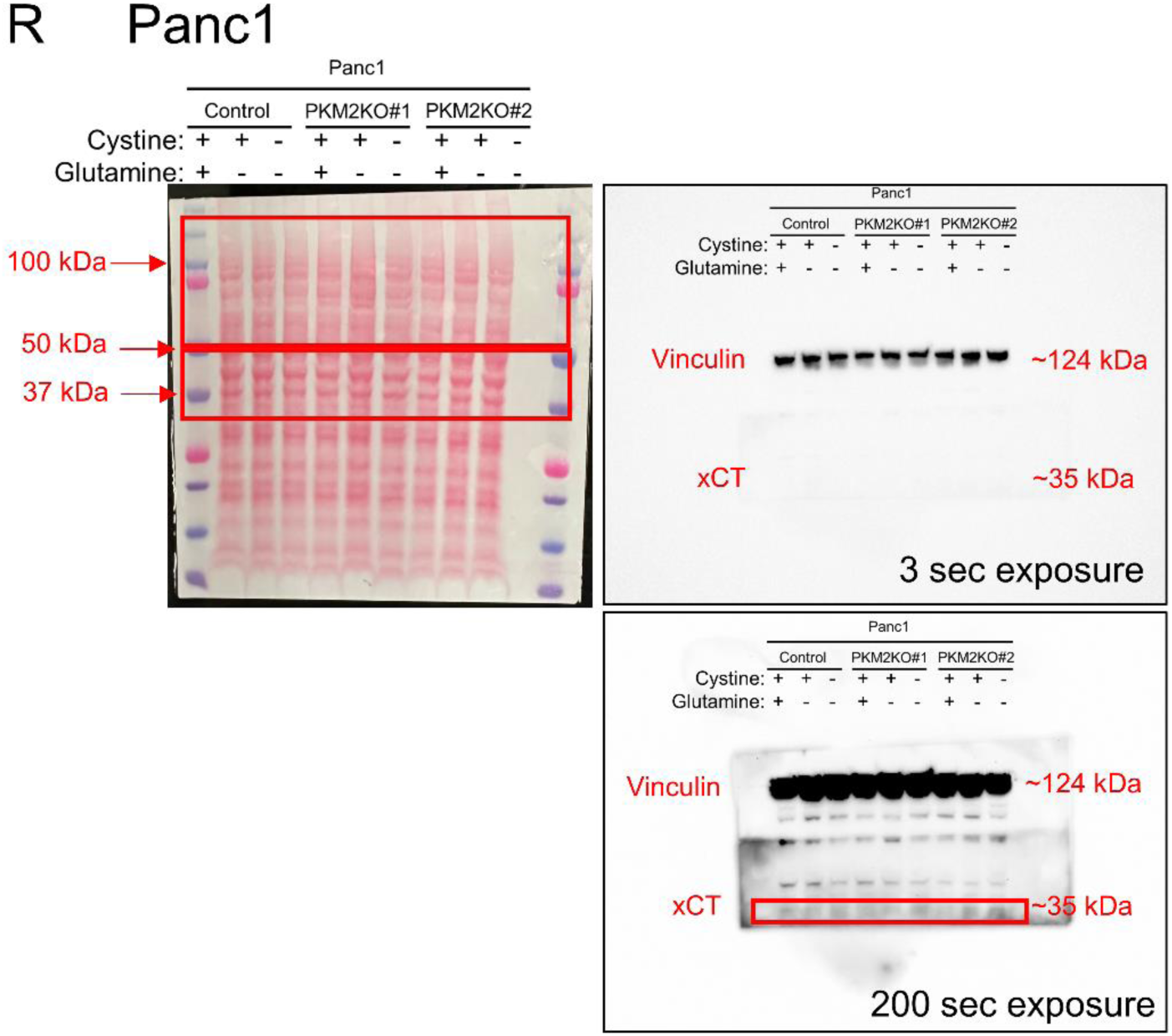
Data in support of Figure 6. **A.** Relative viabilities of AsPC1 control and PKM2KO#1 under 50 μM (+) and 0 μM (-) cystine with (+) or without (-) 250 μM glutamine. **B.** Relative viabilities of AsPC1 control and PKM2KO#1 under 50 μM (+) or 0 μM (-) cystine treated with 5 μM CB-839 (glutaminase inhibitor), supplemented with either 8 mM dimethyl-α-ketoglutarate (α-Kg), 8 mM dimethyl-succinate (Suc), or 32 mM dimethyl-malate (Mal). For **A**-**B**, significance was assessed by two-way ANOVA. \**p*<0.05, \*\**p*<0.01, \*\*\**p*<0.001. Multiple hypothesis correction by Tukey test. **C.** Relative viability of AsPC1 PKM2KO #3 under 50 μM (+) or 0 μM (-) cystine with (+) or without (-) 1 mM glutamine supplemented with either 8 mM dimethyl-α-ketoglutarate (α-Kg), 8 mM dimethyl-succinate (Suc), or 32 mM dimethyl-malate (Mal). Significance was assessed by one-way ANOVA. \**p*<0.05, \*\**p*<0.01, \*\*\**p*<0.001. Multiple hypothesis correction by Sidak test. **D.-F.** Relative viabilities of Panc1 and PKM2KO clones under 50 μM (+) or 0 μM (-) cystine with or without 1 mM glutamine supplemented with either 8 mM DM-αkg, 8 mM DM-Suc, or 32 mM DM-Mal. For **C**-**F**, significance was assessed by one-way ANOVA. \**p*<0.05, \*\**p*<0.01, \*\*\**p*<0.001. Multiple hypothesis correction by Sidak test. **G**-**H**. Relative viabilities of AsPC1 (**A**) and Panc1 (**H**) control and PKM2KO clones under 50 μM (+) or 0 μM (-) cystine supplemented with 8 mM dimethyl-malate (Mal). Significance was assessed by two-way ANOVA. \**p*<0.05, \*\**p*<0.01, \*\*\**p*<0.001. Multiple hypothesis correction by Tukey test. **I-L.** Relative viabilities of AsPC1 and Panc1 (**M-O**) control and PKM2KO clones under 50 μM cystine supplemented with either 8 mM DM-αkg, 8 mM DM-Suc, or 32 mM DM-Mal. Significance was assessed by one-way ANOVA. \**p*<0.05, \*\**p*<0.01, \*\*\**p*<0.001. Multiple hypothesis correction by Sidak test. **P-R.** Full images of Ponceau staining and western blot from cropped image in Figure 6E and 6F respectively. Red boxes indicate cut regions of membrane for primary antibody incubation and cropped portions of immunoblot.

**Supplementary Figure S10.**
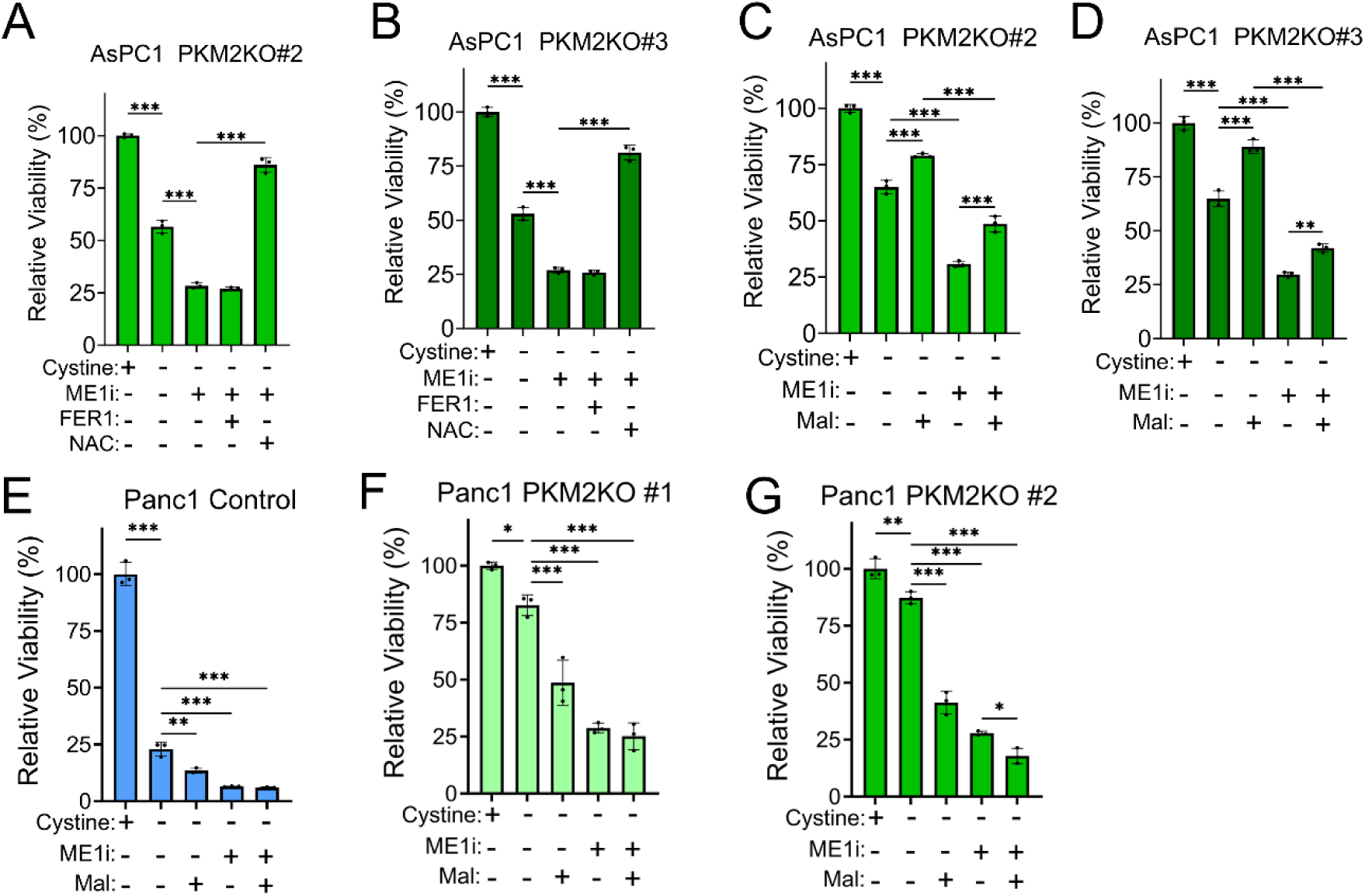
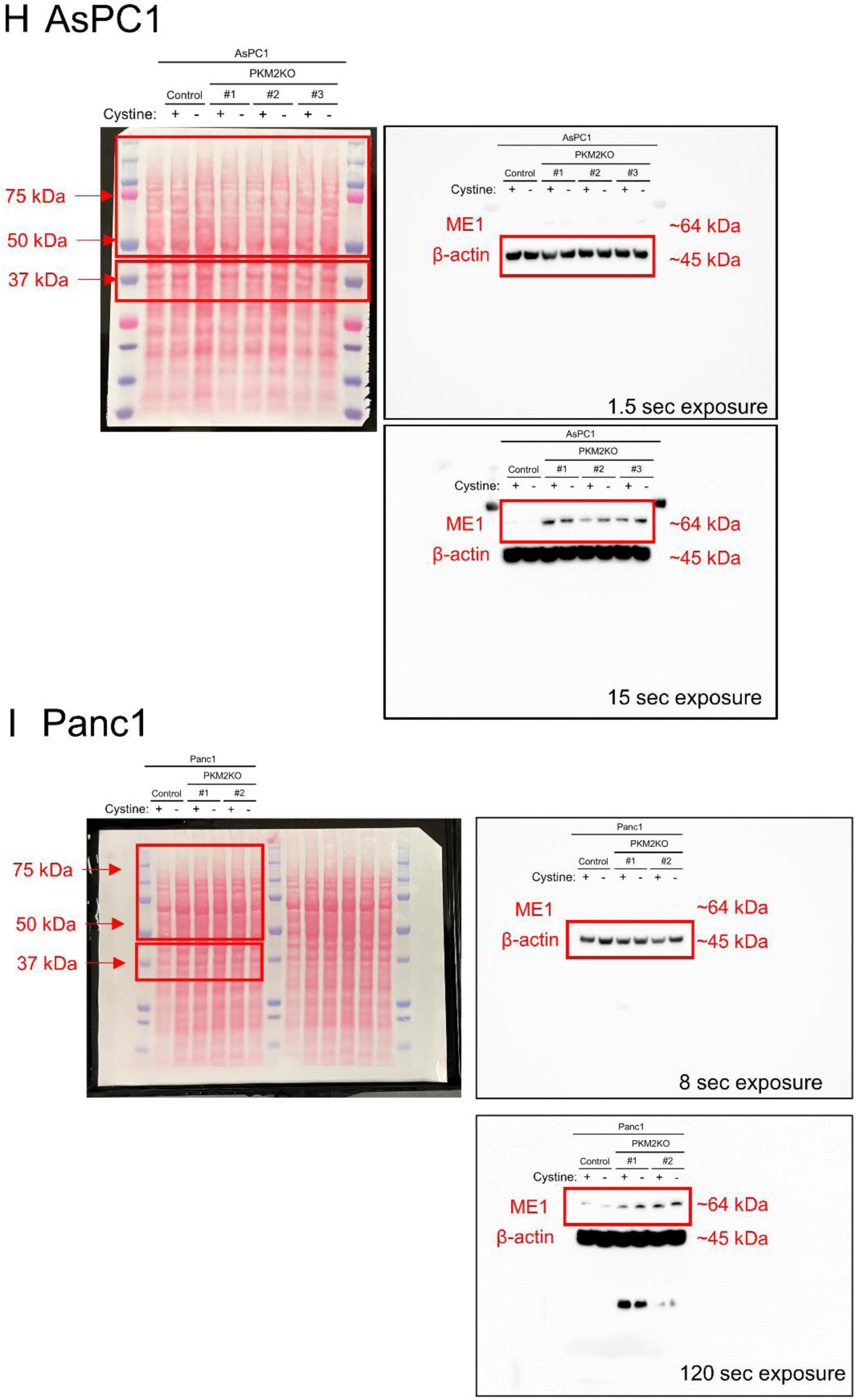
Additional data in support of figure 7. **A**-**B**. Relative viabilities of AsPC1 PKM2KO #2 (**A**) and AsPC1 PKM2KO #3 (**B**) under 0 μM cystine treated with (+) or without (-) 50 μM malic enzyme 1 inhibitor (ME1i) and co-treated with either 5 μM ferrostatin-1 (FER) or 1 mM N-acetylcysteine (NAC). **C**-**G**. Relative viabilities of AsPC1 PKM2KO clone #2 (**A**), AsPC1 PKM2KO #3 (**D**), Panc1 control (**E**), Panc1 PKM2KO #1 (**F**), and PKM2KO #2 (**G**) under 50 μM (+) or 0 μM (-) cystine with or without 50 μM ME1i and 32 mM dimethyl-malate (Mal). For **A**-**G**, significance was assessed by one-way ANOVA. \**p*<0.05, \*\**p*<0.01, \*\*\**p*<0.001. Multiple hypothesis correction by Sidak test. **H.** and **I.** Full images of Ponceau staining and western blot from cropped image in Figure 7A and 7B respectively. Red boxes indicate cut regions of membrane for primary antibody incubation and cropped portions of immunoblot.

